# Imputed Gene Expression Risk Scores: A Functionally Informed Component of Polygenic Risk

**DOI:** 10.1101/2020.12.01.369462

**Authors:** Oliver Pain, Kylie P. Glanville, Saskia Hagenaars, Saskia Selzam, Anna Fürtjes, Jonathan R. I. Coleman, Kaili Rimfeld, Gerome Breen, Lasse Folkersen, Cathryn M. Lewis

## Abstract

**Background:** Integration of functional genomic annotations when estimating polygenic risk scores (PRS) can provide insight into aetiology and improve risk prediction. This study explores the predictive utility of gene expression risk scores (GeRS), calculated using imputed gene expression and transcriptome-wide association study (TWAS) results.

**Methods:** The predictive utility of GeRS was evaluated using 12 neuropsychiatric and anthropometric outcomes measured in two target samples: UK Biobank and the Twins Early Development Study (TEDS). GeRS were calculated based on imputed gene expression levels and TWAS results, using 53 gene expression-genotype panels, termed SNP-weight sets, capturing expression across a range of tissues. We compare the predictive utility of elastic net models containing GeRS within and across SNP-weight sets, and models containing both GeRS and PRS. We estimate the proportion of SNP-based heritability attributable to *cis*-regulated gene expression.

**Results:** GeRS significantly predicted a range of outcomes, with elastic net models combining GeRS across SNP-weight sets improving prediction. GeRS were less predictive than PRS, but models combining GeRS and PRS improved prediction for several outcomes, with relative improvements ranging from 0.3% for Height (*p*=0.023) to 4% for Rheumatoid Arthritis (*p*=5.9×10^-8^). The proportion of SNP-based heritability attributable to *cis*-regulated expression was modest for most outcomes, even when restricting GeRS to colocalised genes.

**Conclusion:** GeRS represent a component of PRS and could be useful for functional stratification of genetic risk. Only in specific circumstances can GeRS substantially improve prediction over PRS alone. Future research considering functional genomic annotations when estimating genetic risk is warranted.

## Introduction

Polygenic risk scores (PRS) are a useful research tool and a promising opportunity for personalised medicine (1). A PRS indicates an individual’s genetic liability for an outcome and is traditionally calculated as the genome-wide association study (GWAS) effect size-weighted sum of alleles (2). The correlation between genetic variants, termed linkage disequilibrium (LD), should be accounted for when estimating PRS. LD-based clumping is often used to obtain LD-independent variants, though more recent methods that estimate the joint effect of variants to account for LD have been shown to improve prediction (3). The predictive utility of PRS can be further increased by incorporating prior probability distributions on causal effect sizes, thereby reducing the signal to noise ratio (4).

A wealth of research has shown enrichment in GWAS of expression quantitative trait loci (eQTLs), variants affecting gene expression (5, 6). The eQTL studies have identified many genetic variants associated with differential gene expression (7, 8). Integration of eQTL and GWAS summary statistics enables inference of gene expression changes associated with the GWAS phenotype, an approach called transcriptome-wide association study (TWAS) (9, 10). TWAS aggregates the effect of genetic associations in a functionally-informed manner to highlight associated up-/down-regulated genes within the context of a specific tissue or developmental stage (11). Due to the functionally informed aggregation of individual genetic effects, TWAS can identify novel associations not previously identified as significant in the corresponding GWAS. This approach has been useful for highlighting plausible candidate genes for experimental follow-up (12).

There has been limited research investigating the predictive utility of PRS that consider the effect of each variant on gene expression. One approach is to split genetic variants into high and low prior groups based on whether they are eQTLs, and then calculate the PRS using a range of mixing parameters to optimally weight the contribution of high prior variants (13). This approach of reweighting eQTL variants improved prediction over functionally agnostic PRS in type 2 diabetes. An alternative approach is to calculate gene-expression risk scores (GeRS), which consider the joint effect of variation on each gene’s expression (12). GeRS are calculated as the sum of predicted expression for an individual weighted by the TWAS-based effect size, analogous to PRS except using predicted expression instead of individual genotypes, and TWAS effect size instead of GWAS effect size. GeRSs were shown to significantly predict schizophrenia, with GeRS derived using prefrontal cortex eQTL data explaining the most variance compared to other individual tissues, but a model containing GeRS based on multiple tissues providing the largest variance explained. However, whether GeRS can improve prediction in combination with PRS was not investigated. A recent study reports that the genetically regulated transcriptome is a component of broader genetic variation, but modelling these sources of variance separately improved out-of-sample prediction (14). This finding suggests that a GeRS will capture a component of PRS, but modelling GeRS and PRS separately will improve prediction

Previous research has shown that GeRS can explain significant variance in schizophrenia, and that modelling variance explained by the genetically regulated transcriptome could improve prediction over models considering the genome alone. However, GeRS have only been applied to schizophrenia, and no previous study has combined GeRS with PRS. In this study, we evaluate the predictive utility of GeRS calculated using the TWAS-based approach with eQTL data from a range of tissues. We apply the method to a range of quantitative traits and binary disorders in two samples, UK Biobank (UKB) (15) and the Twins Early Development Study (TEDS) (16). Furthermore, we evaluate whether GeRS provide novel information over PRS and explore the effect of stratifying genes by colocalization estimates of pleiotropy and tissue specificity of eQTL effects.

## Methods

### UK Biobank (UKB)

UKB is a prospective cohort study that recruited >500,000 individuals aged between 40-69 years across the United Kingdom (15). The UKB received ethical approval from the North West - Haydock Research Ethics Committee (reference 16/NW/0274).

#### Genetic data

UKB released imputed dosage data for 488,377 individuals and ~96 million variants, generated using IMPUTE4 software (15) with the Haplotype Reference Consortium reference panel (17) and the UK10K Consortium reference panel (18). This study retained individuals that were of European ancestry based on 4-means clustering on the first 2 principal components provided by the UKB, had congruent genetic and self-reported sex, passed quality assurance tests by UKB, and removed related individuals (>3^rd^ degree relative, KING threshold > 0.044) using relatedness kinship (KING) estimates provided by the UKB (15). The imputed dosages were converted to hard-call format for all variants.

#### Phenotype data

Eight UKB phenotypes were analysed. Six phenotypes were binary: Depression, Type 2 Diabetes (T2D), Coronary Artery Disease (CAD), Inflammatory Bowel Disease (IBD), and Rheumatoid arthritis (RheuArth). Three phenotypes were continuous: Intelligence, Height, and Body Mass Index (BMI). Further information regarding outcome definitions can be found in the Supplementary Material.

Analysis was performed on a subset of ~50,000 UKB participants for each outcome to reduce the computational burden of the analysis whilst maintaining sufficient power to perform downstream analyses. For each continuous trait (Intelligence, Height, BMI), a random sample was selected. For disease traits, all cases were included, except for Depression and CAD where a random sample of 25,000 cases was selected. Controls were randomly selected to obtain a total sample size of 50,000. Sample sizes for each phenotype after genotype data quality control are shown in Table 1.

**Table 1.** Sample size for each target sample phenotype

| **UKB**<br>**Phenotype** | **Description** | **Total sample**<br>**size** | **No. of**<br>**controls** | **No. of cases** |
| --- | --- | --- | --- | --- |
| Depression | Major depression | 49995 | 24999 | 24996 |
| Intelligence | Fluid intelligence | 49998 | NA | NA |
| BMI | Body Mass Index | 49993 | NA | NA |
| Height | Height | 49993 | NA | NA |
| T2D | Type-2 Diabetes | 49990 | 35102 | 14888 |
| CAD | Coronary Artery Disease | 49991 | 24998 | 24993 |
| IBD | Inflammatory Bowel<br>Disease | 50000 | 46539 | 3461 |
| RheuArth | Rheumatoid Arthritis | 50000 | 46592 | 3408 |
| **TEDS**<br>**Phenotype** |  |  |  |  |
| GCSE | Mean GCSE scores | 7296 | NA | NA |
| ADHD | ADHD symptoms | 7880 | NA | NA |
| BMI21 | Body Mass Index at age 21 | 5220 | NA | NA |
| Height21 | Height at age 21 | 5455 | NA | NA |

### TEDS

The Twins Early Development Study (TEDS) is a population-based longitudinal study of twins born in England and Wales between 1994 and 1996 (16). Ethical approval for TEDS has been provided by the King’s College London ethics committee (reference: 05/Q0706/228). Parental and/or self-consent was obtained before data collection. For this study, one individual from each twin pair was removed to retain only unrelated individuals.

#### Genetic data

As previously described (19), TEDS genotype data underwent stringent quality control prior to imputation via the Sanger Imputation server using the Haplotype Reference Consortium reference data (17). Imputed genotype dosages were converted to hard-call format using a hard call threshold of 0.9, with variants for each individual set to missing if no genotype had a probability of > 0.9. Variants with an INFO score < 0.4, MAF < 0.001, missingness > 0.05 or Hardy-Weinberg equilibrium p-value < 1×10^-6^ were removed.

#### Phenotypic data

This study used four continuous phenotypes within TEDS: Height, Body Mass Index (BMI), Educational Achievement (GCSE), and Attention Deficit Hyperactivity Disorder (ADHD) symptom score (Table 1). Further information regarding the phenotype definitions can be found in the supplementary material and a previous study (20).

### Genotype-based Scoring

Gene expression risk scores (GeRS) and polygenic risk scores (PRS) were calculated within a reference-standardised framework, whereby the resulting PRS and GeRS are not influenced by target sample specific properties including availability of variants and measurements of LD and allele frequency. This is achieved by using a common set of typically well imputed variants (HapMap3) and using reference genetic data (European 1KG Phase 3) to estimate LD and allele frequencies. Lastly, all genotype-based scores are scaled and centred based on the mean and standard deviation of scores in the reference sample. This reference-standardised approach and its merits have been described previously (3).

A schematic representation of calculating GeRS is shown in Figure 1.

**Figure 1.**
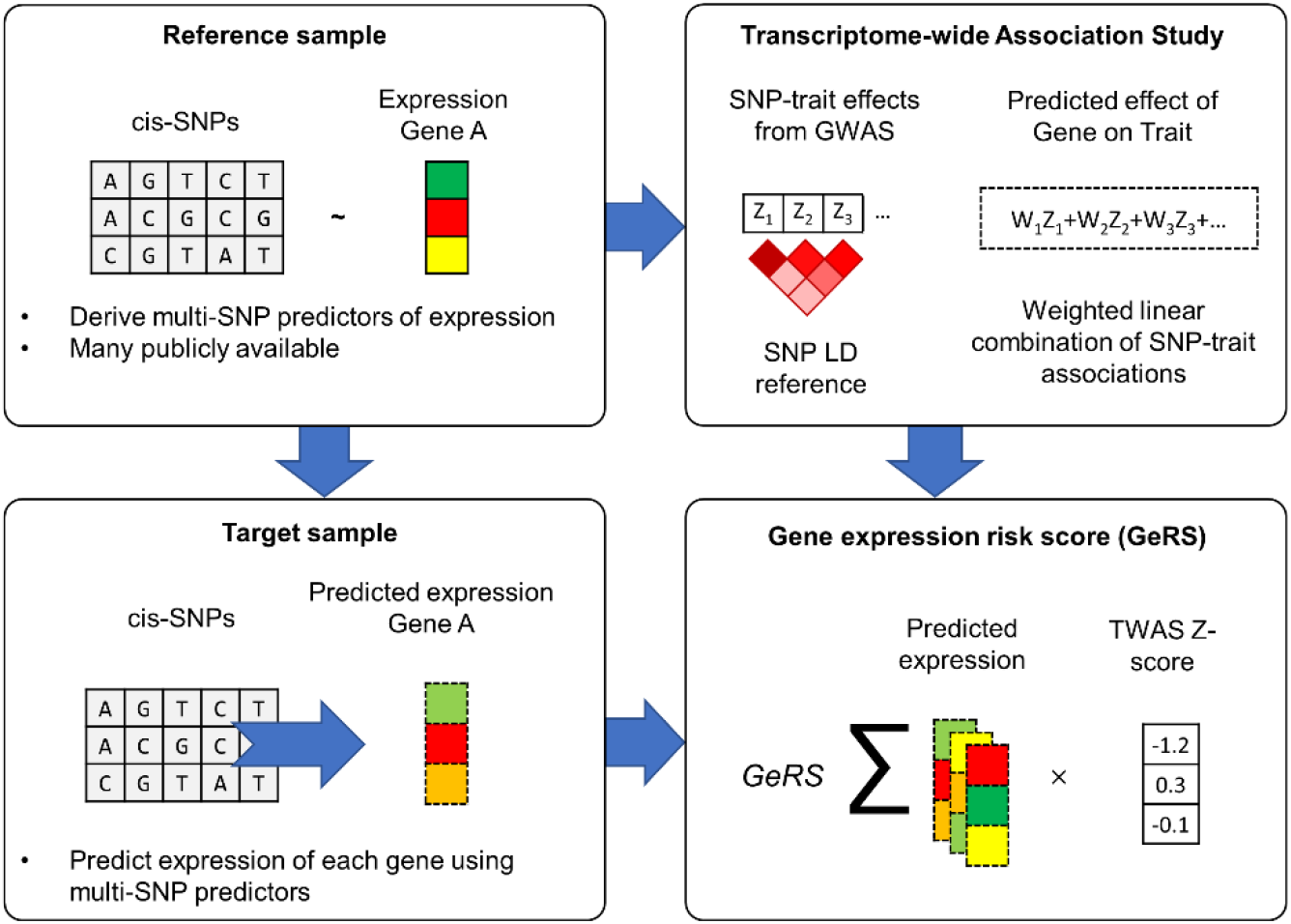
Schematic representation of GeRS Calculation. The top left panel describes the process of deriving SNP-weights predicting gene expression in a sample of individuals with both genotype and gene expression measured (e.g. Genotype-tissue expression consortium, GTEx). These SNP-weights can be used to perform TWAS whereby SNP-weights are integrated with GWAS summary statistics to infer gene expression associations with the GWAS trait (upper right panel). The SNP-weights can also be used to predict gene expression levels in a target sample with only genotype data available (lower left panel). Finally, GeRS can be calculated in the target sample by combining the level of predicted expression in each individual weighted by the TWAS effect size (lower right panel). This figure has been adapted from Gusev et al. 2016 (9).

#### GWAS summary statistics

GWAS summary statistics were identified for phenotypes the same as or similar as possible to the UKB and TEDS phenotypes (descriptive statistics in Table S1), excluding GWAS with documented sample overlap with the target samples. GWAS summary statistics were formatted using the LD-Score Regression munge_sumstats.py script (see Web Resources) with default settings (listed in the Supplementary Material) except the minimum INFO score was set to 0.6.

#### Transcriptome-wide association study (TWAS)

FUSION software (9) was used to integrate GWAS summary statistics with precomputed SNP-weights of gene expression to infer differential gene expression associated with the GWAS-phenotype. The term *SNP-weight* refers to a multi-SNP-based predictor of a gene’s expression. SNP-weights used in this study were derived using gene expression data from a range of distinct tissues and European-ancestry adulthood samples, downloaded from the FUSION website (See URLs). The weights pertained to five RNA reference samples: (i) the Genotype-Tissue Expression (GTEx) Consortium (Version 7)(7), measuring gene expression across 48 tissues, including brain regions, blood and peripheral tissues, (ii) The CommonMind Consortium (CMC)(8), measuring expression and differential splicing in the dorsolateral prefrontal cortex, (iii) The Netherlands Twins Register (NTR)(21) and (iv) Young Finns Study (YFS)(9), which both provide information on blood tissue gene expression, and (v) Metabolic Syndrome in Men (METSIM)(9), assessing adipose tissue expression. The SNP-weights obtained from a given sample and tissue (e.g. GTEx thyroid, NTR peripheral blood) are referred to as *SNP-weight sets*. Characteristics for the 53 SNP-weight sets used are available in Table S2. The SNP-weights include 260,598 *features* (SNP-weight set and gene pairs), capturing expression of 26,434 unique genes (protein-coding and non-protein coding). The number of features that could be reliably imputed for each GWAS is shown in Table S3. TWAS was performed using default settings in FUSION and LD estimates from the European subset of the 1KG Phase 3 reference sample (N=503).

Colocalization analysis tests whether the association between a genetic locus and two or more traits is driven by the same causal variant, or whether the association for each trait is driven by different causal variants that are in LD. Colocalization was performed using the coloc R package (22), implemented within the FUSION software, to estimate the posterior probability that the GWAS phenotype and gene’s expression share a single causal variant, termed PP4. A coloc *p*-value threshold of 0.05 was used, to perform colocalization for all features with a TWAS *p*-value < 0.05.

#### Predicting expression in target samples

The *cis*-heritable component of expression for each gene was imputed in each target sample using the same gene expression SNP-weights described above, and target sample genotype data. Predicted expression levels are calculated as,

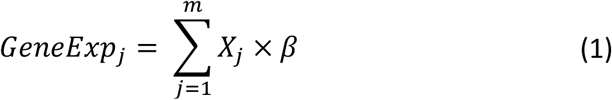

where the predicted level of expression (*GeneExp*) for an individual is the number of effect alleles carried by the individual (*X_j_*) weighted by the effect of each variant on gene expression as estimated from penalised regression model (β), across *m* variants. This was implemented using the FUSION script ‘make_score.R’ to convert the TWAS SNP-weights into PLINK score file format, and then using PLINK to carry out the scoring in the target sample. Predicted expression levels are then centred and scaled based on the mean and standard deviation of the predicted expression in the 1KG Phase 3 European reference sample.

#### Gene Expression Risk Scoring

Gene expression risk scores (GeRS) were calculated as

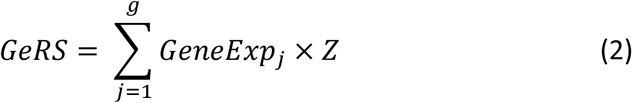

where the *GeRS* of an individual is equal to the TWAS effect size (Z)-weighted sum of the individual’s predicted expression (*GeneExp_j_*), at *g* genes. GeRS were calculated for each SNP-weight set separately, meaning 53 GeRS for each GWAS/TWAS phenotype were generated. To remove genes with highly correlated predicted expression due to LD, genes were ranked by TWAS p-value and clumping was performed to remove genes with a predicted expression *R*^2^ > 0.9 within 500kb of the lead gene boundaries. Within the MHC region, the single most significant gene was retained due to long range and complex LD structures. Predicted expression estimates used for clumping were estimated in the European 1KG Phase 3 reference. A range of nested *p*-value thresholds were used to select genes considered in the GeRS: 1, 5×10^-1^, 1×10^-1^, 5×10^-2^, 1×10^-2^, 1×10^-3^, 1×10^-4^, 1×10^-5^ and 1×10^-6^. Scripts used to perform gene expression risk scoring can be found on the GenoPred website (see URLs).

In addition, we evaluate the predictive utility of GeRS restricted to genes with evidence of colocalization with the outcome (PP4 > 0.8), and GeRS restricted to genes showing tissue specific expression. Tissue-specific GeRS were derived by only considering genes that were either not significantly heritable in blood SNP-weight sets (GTEx Whole blood, YFS or NTR), or genes whose predicted expression was uncorrelated with the corresponding feature in the blood SNP-weight sets (*R*^2^ < 0.01). This approach is congruent with a previous study identifying tissue specific eQTL effects prior to risk scoring (23). Blood-specific features were identified using the same criteria but comparing predicted expression across all non-blood SNP-weight sets. The number of tissue-specific features for each SNP-weight set are listed in Table S2.

#### Polygenic Risk Scores (PRS)

Polygenic scoring was carried out using the traditional *p*-value thresholding and LD-based clumping approach (pT+clump), and a more recent method, PRScs (24), which models LD between genetic variants and applies shrinkage parameters to avoid overfitting. PRScs has been previously reported to out-perform other polygenic scoring methods (3). pT+clump was performed using an *R^2^* threshold of 0.1 and window of 250kb. Within the MHC region (28-34Mb on chromosome 6), the pT+clump method retains only the single most significant variant due to long range of complex LD in this region. A range of p-value thresholds were used to select variants: 1×10^-8^, 1×10^-6^, 1×10^-4^, 1×10^-2^, 0.1, 0.2, 0.3, 0.4, 0.5 and 1. PRScs was performed using a range of global shrinkage parameters (1×10^-6^,1×10^-4^,1×10^-2^ and 1) and the fully Bayesian mode, which estimates the optimal shrinkage parameter. Analogous to the GeRS, only HapMap3 variants were considered during polygenic scoring, and the European subset of the 1KG Phase 3 reference was used to estimate LD.

As a sensitivity analysis, pT+clump PRS were also calculated using only variants within 500kb of genes used in the TWAS, thereby restricting the PRS to the same variants within the gene expression SNP-weights and highlighting the effect of reweighting genetic variants by their effect on gene expression.

Furthermore, pT+clump PRS for Rheumatoid Arthritis were also calculated without restricting to a single variant in the MHC region to gain insight into difference between PRS and GeRS prediction for this outcome.

### Evaluating predictive utility of GeRS

Prediction accuracy was evaluated as the Pearson correlation between the observed and predicted phenotype outcomes. Correlation was used as the main test statistic as it is applicable for both binary and continuous outcomes and standard errors are easily computed. Correlations can be easily converted to other test statistics such as *R^2^* (observed or liability) and area under the curve (AUC) (equations 8 and 11 in (25)), with relative performance of each method remaining unchanged.

Logistic regression was used for predicting binary outcomes, and linear regression was used for predicting continuous outcomes. If the model contained only one predictor, a generalized linear model was used. If the model contained more than one predictor (e.g. GeRS for each *p*-value threshold), an elastic net model was applied to avoid overfitting due to the inclusion of multiple correlated predictors (26).

#### Elastic net modelling

Previous research has shown that modelling multiple PRS based on a range of parameters (*p*-value thresholds or shrinkage parameters) optimises prediction out-of-sample (3). Therefore, elastic net models were derived using pT+clump PRS across *p*-value thresholds, or PRScs scores across global shrinkage parameters. Furthermore, elastic net models were derived for GeRSs across a range of *p*-value thresholds and SNP-weight sets to evaluate the effect of modelling multiple GeRS simultaneously.

A nested cross validation procedure (27) was used to estimate the predictive utility of elastic net models (Figure 2), where hyperparameter selection is performed using inner 10-fold cross validation, while an outer 5-fold cross-validation computes an unbiased estimate of the predictive utility of the model with the inner cross validation based hyperparameter tuning. This approach avoids overfitting whilst providing modelling predictions for the full sample. The inner 10-fold cross validation for hyperparameter optimisation was carried out using the ‘caret’ R package.

**Figure 2.**
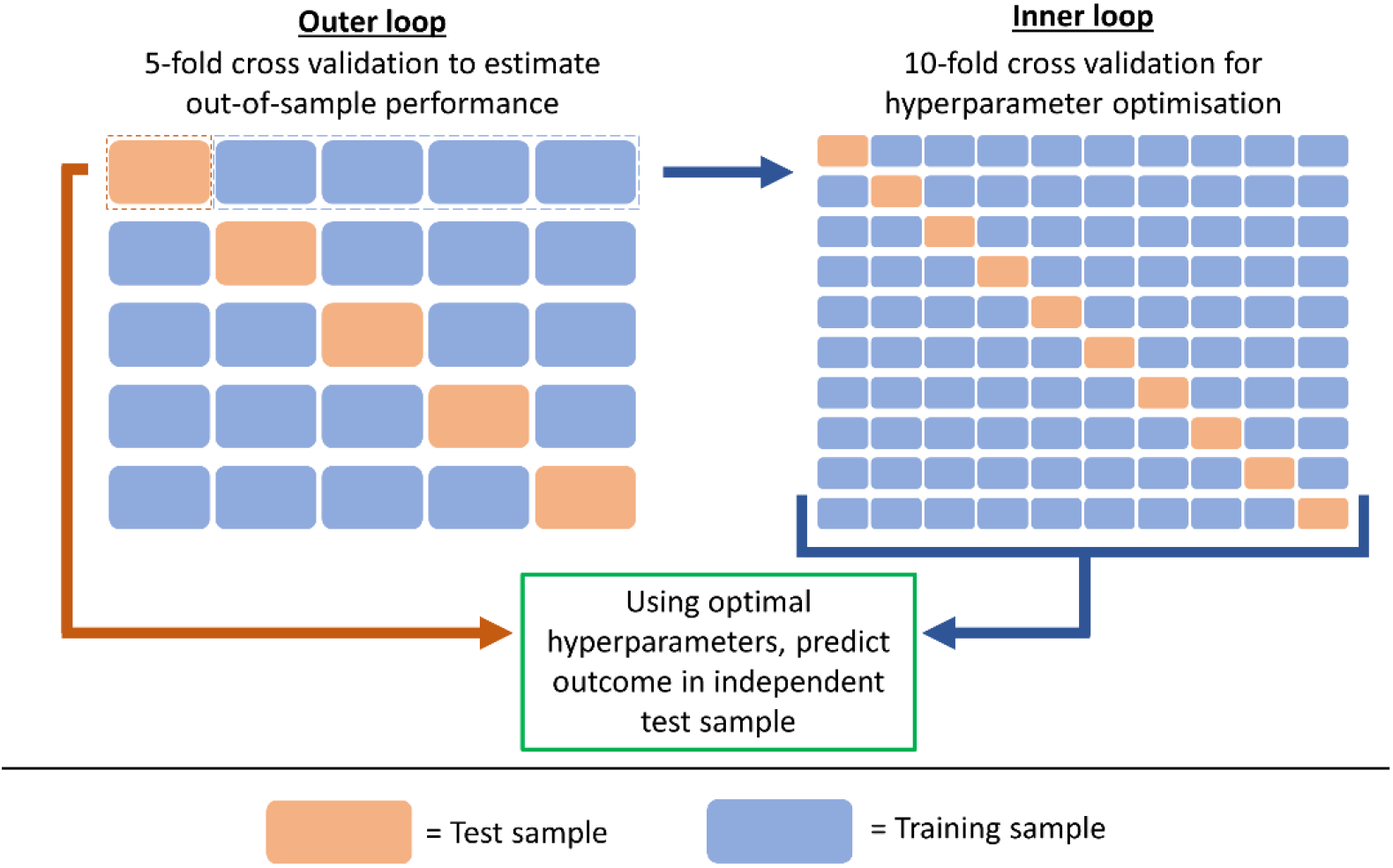
Schematic representation of nested cross validation procedure. The outer loop splits the sample into 5 parts, 4 parts are used as a training sample for hyperparameter optimisation, and the resulting model is then used to predict the outcome in the remaining part (test sample). This process is repeated until predictions are available for all parts of the sample. Hyperparameter optimisation is carried out within the inner loop, which consists of 10-fold cross validation.

The correlation between observed and predicted values of each model were compared using William’s test (also known as the Hotelling-Williams test) (28) as implemented by the ‘psych’ R package’s ‘paired.r’ function, with the correlation between model predictions of each method specified to account for their non-independence. A two-sided test was used when calculating *p*-values.

#### Estimating variance explained by cis-heritable expression

A schematic representation of this analysis is in Figure S1. To estimate the proportion of SNP-based heritability explained by *cis*-regulated expression, we used AVENGEME to estimate SNP-based heritability of each phenotype in the target sample based on pT+clump PRS associations across *p*-value thresholds, and the phenotypic variance explained by *cis*-regulated expression (GE-based heritability) based on the GeRS associations across *p*-value thresholds. To estimate the association with GeRS at each *p*-value threshold we used predictions from elastic net models containing GeRS across all SNP-weight sets for a given *p*-value threshold. The proportion of SNP-based heritability explained by *cis*-heritable expression was then calculated as GE-based heritability divided by the SNP-based heritability. AVENGEME also estimates the fraction of non-causal variants. AVENGEME has been previously used to estimate the proportion of SNP-based heritability attributable to *cis*-regulated gene expression based on GeRS associations, acknowledging that the estimate will be inflated due to LD causing gene expression SNP-weights to tag other causal mechanisms, such as variants affecting protein structure and function (12). As a sensitivity analysis, we estimated the GE-based heritability using GeRS restricted to genes with colocalization PP4 > 0.8 to remove genes which do not colocalise. For the GeRS analysis, the ‘nsnp’ variable in AVENGEME, indicating the number of independent markers in the score was set to the number of LD independent markers in the TWAS gene stratified PRS.

## Results

### Predictive utility of GeRS

For the six disorders and six quantitative traits analysed in UK Biobank and TEDS, the GeRS calculated were significantly correlated with each phenotype. GeRS were most predictive of Height in TEDS using the GTEx Nerve Tibial SNP-weight set with a correlation between predicted and observed values of 0.22 (SE=0.01, *p*-value= 6.8×10^-61^). The predictive utility of GeRS typically increased as more relaxed *p*-value thresholds were used to select genes (Figures 3, S2-S3). The predictive utility of GeRS for outcomes available in both UKB and TEDS, Height and BMI, were broadly consistent.

**Figure 3.**
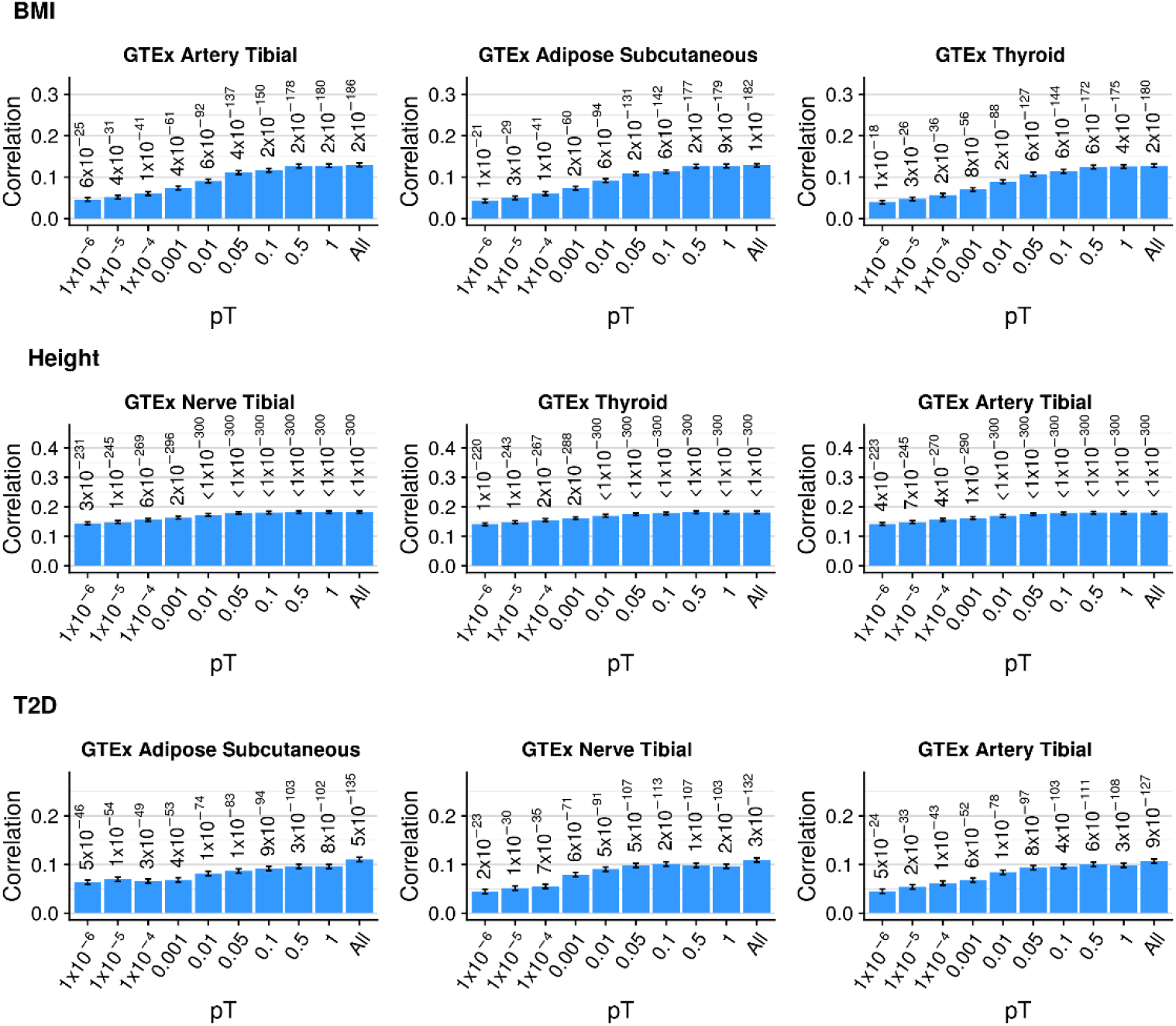
Correlation between GeRS for three outcomes in UKB across p-value thresholds. The all bar indicates the correlation between observed and predicted values from an elastic net model including all p-value thresholds. Error bars represent the standard error of the correlation. GeRS are based on a single SNP-weight sets. Figure only shows results for the three SNP-weights with the strongest correlation between predicted and observed values. Values above bars are p-values indicating whether the correlation is significantly different from zero.

Combining GeRS across *p*-value thresholds in an elastic net model significantly improved prediction over the single best GeRS *p*-value threshold for all outcomes in UKB except Depression and IBD (Figure 4A, Table S4). The largest improvement in prediction when modelling multiple *p*-value thresholds was for T2D in UKB (23.6% improvement, *p*-value=2.2×10^-28^). Modelling GeRS across multiple *p*-value thresholds did not improve prediction for any outcome in the TEDS sample, and led to a significant decrease in prediction for GCSE (6.1% reduction, *p*-value=1.9×10^-3^) (Figure S4A, Table S4).

**Figure 4.**
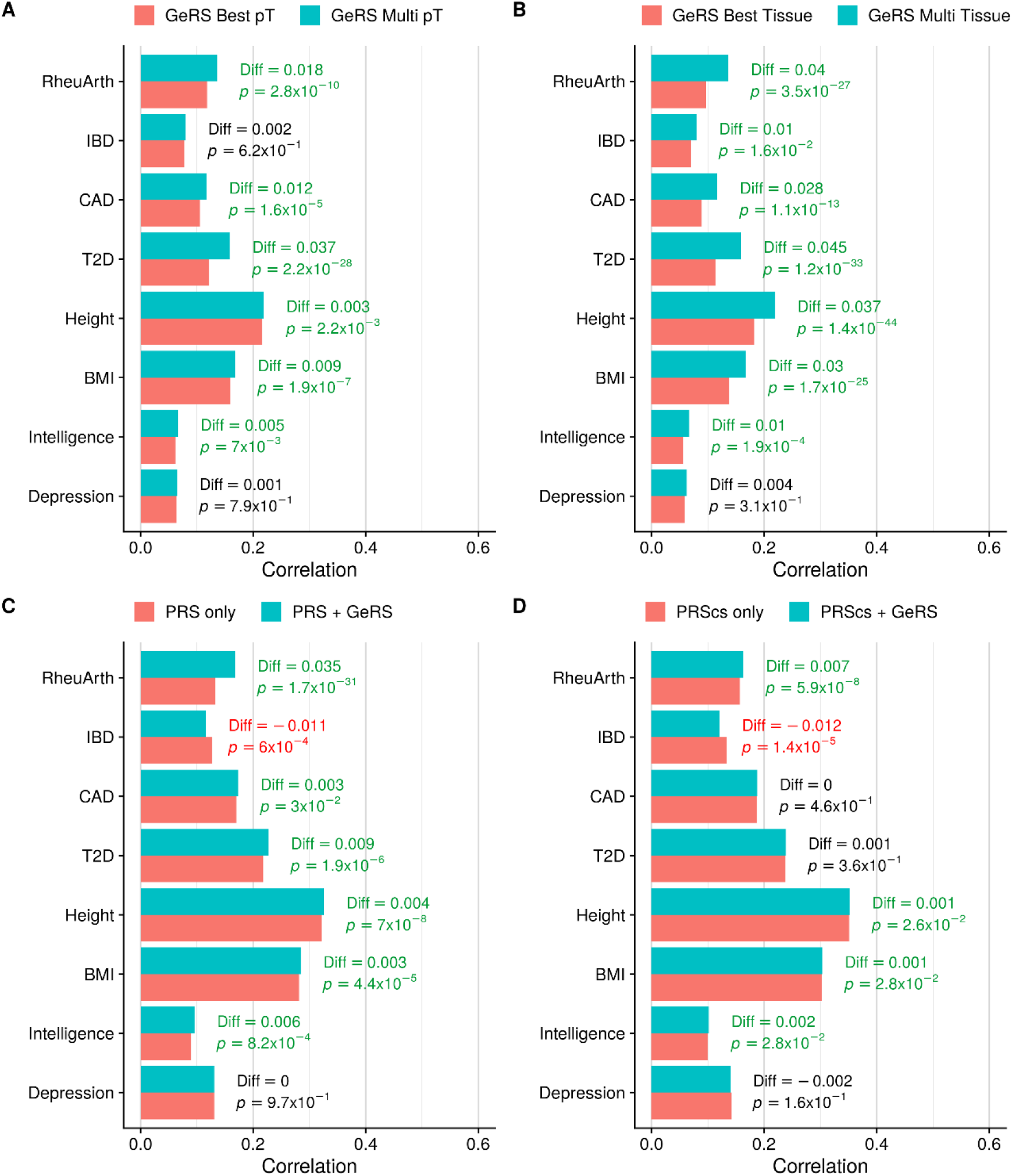
Comparing the predictive utility of GeRS and PRS in UKB. A) Compares the predictive utility of models containing GeRS across SNP-weight sets based on the single best p-value threshold and models containing GeRS across all p-value thresholds. B) Compares the predictive utility of models containing GeRS across p-value thresholds based on the single best SNP-weight set and models containing GeRS based on all SNP-weight sets. C) Compares the predictive utility of models containing PRS and models containing GeRS and PRS. D) Compares the predictive utility of models containing models also containing PRS derived using PRScs, and models also containing GeRS. Values on the right of each bar indicate the absolute difference in predicted-observed correlation between the full and nested model. Values on the right are coloured in green to indicate a significant increase in prediction, and red to indicate a significant decrease in prediction.

Modelling GeRS derived using multiple SNP-weights significantly improved prediction over any single SNP-weight set for all outcomes except Depression in UKB, and GCSE and ADHD symptoms in TEDS (Figure 4B and S4B, Table S4). Significant relative improvements provided by modelling GeRS from multiple SNP-weight sets varied from 7.1% (*p*-value = 2.4×10^-2^) for Height in TEDS to 29% (*p*-value = 3.5×10^-27^) for RheuArth in UKB.

### Comparison of SNP-weight sets

The predictive utility of GeRS derived using each SNP-weight set separately is shown in Figures S5-S6. Often the most predictive GeRS were derived using SNP-weight sets capturing expression in tissues previously implicated for the outcome, such as CMC DLFPC for Depression and BMI in UKB. However, the predictive utility of GeRS showed a strong relationship with the size of the sample used to derive the SNP-weights (*r_pearson_*=0.15 in UKB), and the number of features within the SNP-weight sets (*r_pearson_*=0.29 in UKB) (Figures 5 and S7). When fitting both the SNP-weight set sample size and number of features in a joint model, the effect of sample size was no longer significant. After correcting for the number of features in each SNP-weight set, the most predictive SNP-weight set varied for most outcomes (Figures S8-S9). For example, the most predictive SNP-weight set for Depression was GTEx Thyroid but changed to CMC DLPFC after accounting for the number of features within each SNP-weight set. The CMC DLPFC Splicing SNP-weight set was often the least predictive after correcting for the number of features due to features often capturing multiple splice variants for a given gene which are therefore highly redundant.

**Figure 5.**
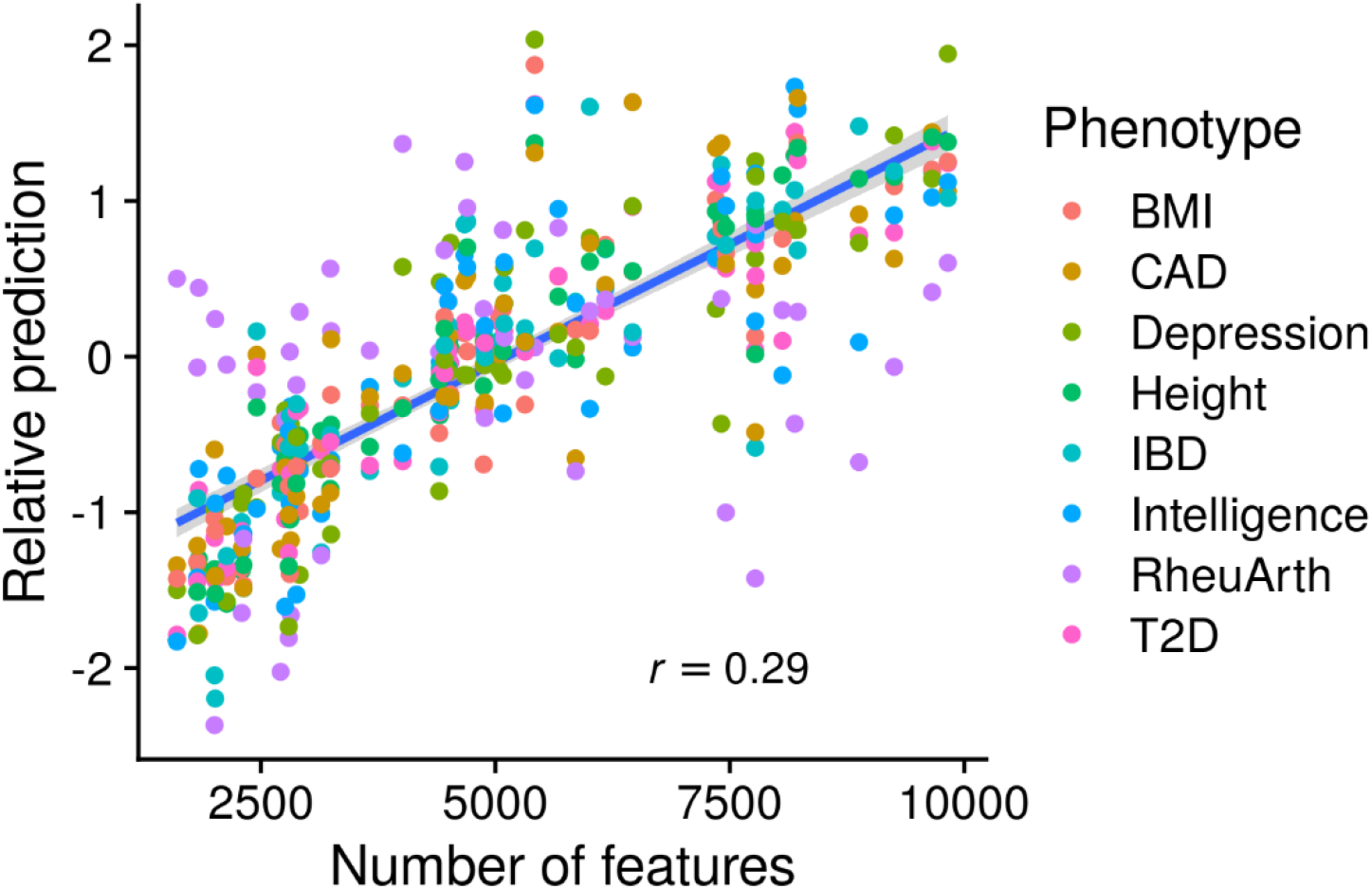
Relationship between the predictive utility of GeRS and the number of features within each SNP-weight set in UKB. The y-axis shows the correlation between observed and predicted values, standardised within each outcome.

### Stratifying by colocalization and tissue specificity

TWAS associations can be driven by the same causal variant driving the association with both gene expression and the phenotype (vertical or horizontal pleiotropy), or the associations can be driven by linkage disequilibrium between different causal variants affecting each outcome. As a result, TWAS associations do not necessarily indicate that the *observed* differential expression of a gene is associated with the outcome. Colocalization estimates of whether both gene expression and the outcome are affected by the same causal variant (PP4), were used to determine whether restricting GeRS to colocalised associations altered the predictive utility of GeRS. We found GeRS restricted to colocalised genes (PP4 > 0.8) had reduced predictive utility compared to unrestricted GeRS (Figures 6, S10-S12).

**Figure 6.**
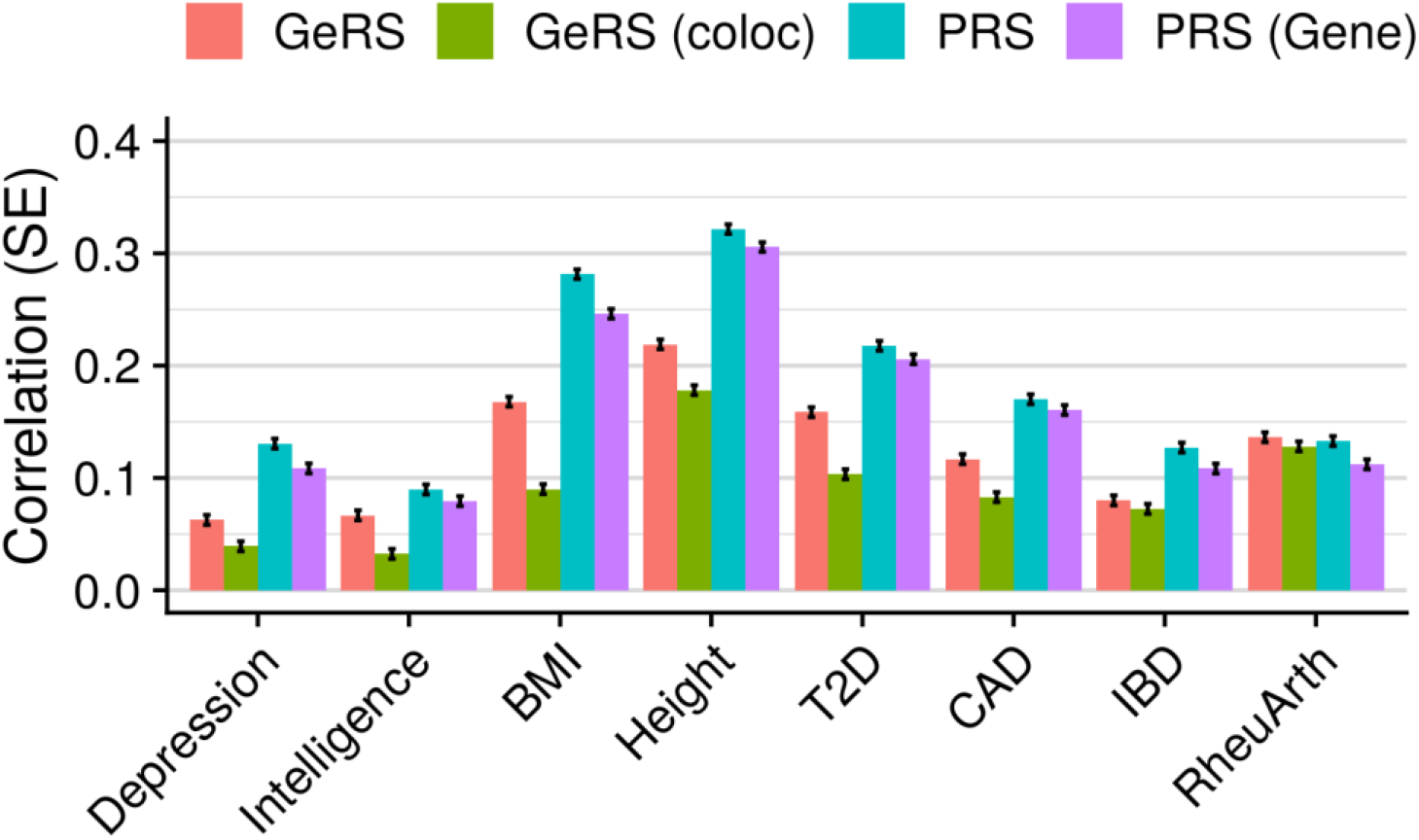
Shows the correlation between predicted and observed values in UKB for models. GeRS = All SNP-weight set GeRS; GeRS (coloc) = All SNP-weight set GeRS restricted to genes with a colocalization PP4 > 0.8; PRS = Genome-wide PRS; PRS (Gene) = PRS restricted to gene regions considered by GeRS.

Cis-eQTL effects are moderately correlated across tissues (7), meaning GeRS for a given SNP-weight set will capture variance attributable to other tissues. To explore the predictive utility of tissue-specific GeRS, we restricted GeRS to genes either not significantly heritable in blood SNP-weight sets, or genes whose predicted expression was uncorrelated with the corresponding feature in the blood SNP-weight sets. We found restricting GeRS to tissue specific features reduced the predictive utility of GeRS based on individual SNP-weight sets, but the predictive utility of models including all SNP-weight sets did not change substantially (Figures S10-S11).

**Figure 7.**
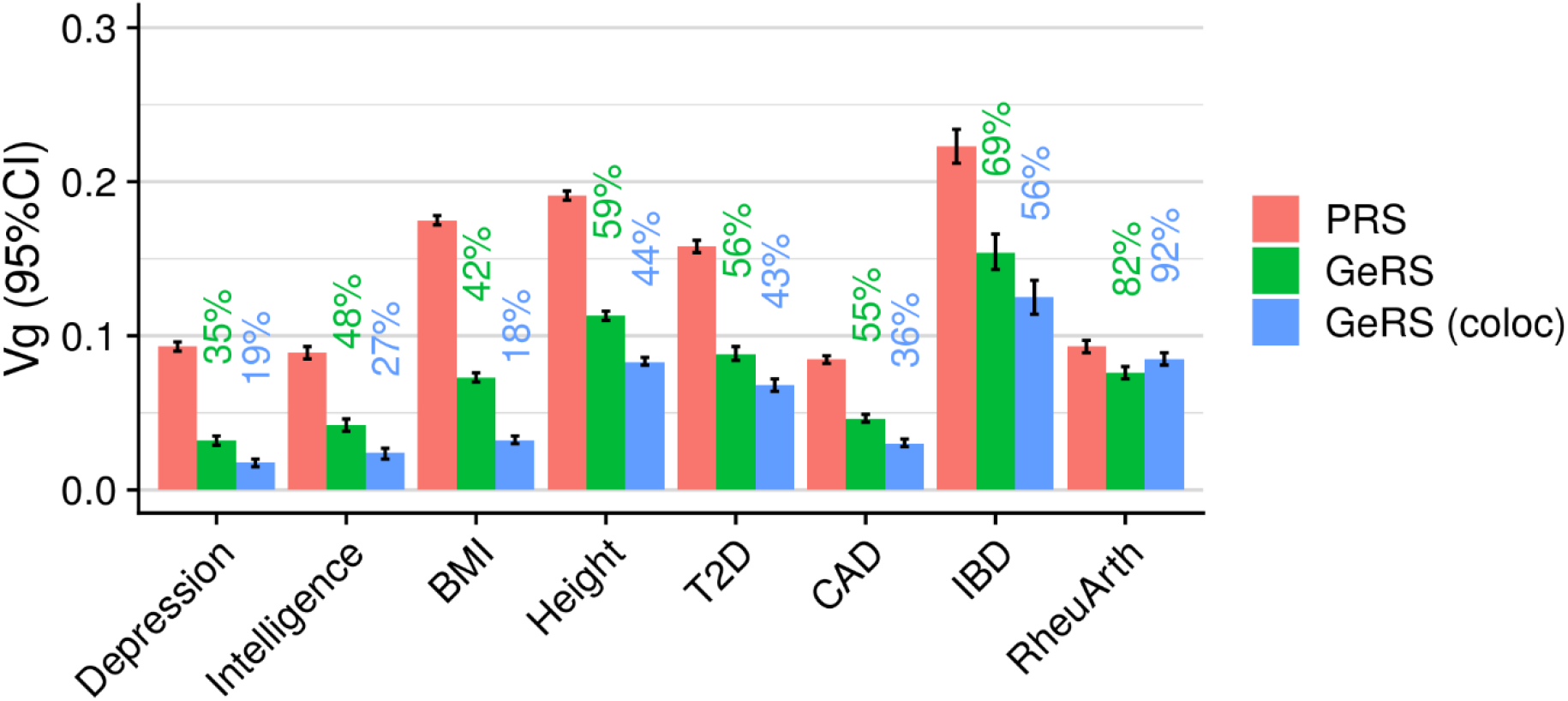
Estimates of SNP-based heritability and GE-based heritability for outcomes in UKB. PRS indicates the SNP-heritability as estimated using PRS association results in AVENGEME. GeRS indicates the GE-based heritability as estimated using GeRS association results in AVENGEME. GeRS (coloc) indicates the GE-based heritability as estimated using GeRS when restricted to genes with colocalization PP4 > 0.8. The value above each bar indicates the proportion of SNP-based heritability accounted for by cis-regulated expression (Green=GeRS/PRS, Blue=GeRS (coloc)/PRS)).

### Comparison of GeRS to PRS

Compared to PRS-only models, models containing PRS and multi-SNP-weight set GeRS provided statistically significant improvements in prediction for all outcomes in UKB except Depression and IBD (Figure 4C, Table S4). Inclusion of GeRS did not significantly improve prediction over PRS-only models for any outcome in TEDS (Figure S4C, Table S4). Statistically significant relative improvements varied from 1% (*p*-value=4.4×10^-5^, correlation increased from 0.281 to 0.284) for BMI in UKB to 20.8% for RheuArth in UKB (*p*-value=1.7×10^-31^, correlation increased from 0.133 to 0.168). Inclusion of GeRS significantly decreased the correlation between observed and predicted values for IBD in UKB (−9.6%, *p*-value=6×10^-4^). We then explored whether GeRS improve prediction over PRS derived using PRScs, which models LD to estimate the joint effect of nearby variants, as opposed to LD-based clumping which removes variants in LD. When comparing GeRS to PRScs scores, the improvement in prediction provided by GeRS was attenuated for all outcomes, although statistically significant relative improvements remained when including GeRS for RheuArth (4%), Height (0.3%), BMI (0.4%) and Intelligence (2.5%) in UKB (Figure 4C, Table S4).

A distinction between the pT+clump PRS and GeRS, is how they handle the MHC region. The pT+clump PRS retain a single variant in the MHC region. In contrast, GeRS retain a single gene in the MHC region, which considers information across multiple variants. Given the large genetic effects in the MHC region for RheuArth, we performed a sensitivity analysis to explore whether the approach of retaining only the single variant in the MHC region is responsible for the improved prediction when including GeRS. The analysis showed that inclusion of GeRS still significantly improved prediction of RheuArth over PRS alone (*p*-value=6.40×10^-11^), though the relative improvement was attenuated from 20.8% to 8.4%.

When comparing the predictive utility of all SNP-weight set GeRS to PRS, we found the proportion of PRS-phenotype correlation that GeRS can explain (*r*_GeRS_/*r*_PRS_) was between 44.6% for BMI in TEDS, and 102.6% for RheuArth in UKB (Figure 6 and S12, Table S5). When restricting GeRS to colocalised genes (PP4 > 0.8), the proportion of PRS-phenotype correlation that GeRS can explain reduced to between −2.8% for ADHD in TEDS and 96.4% for RheuArth in UKB. The predictive utility of PRS stratified to include only variants within gene boundaries was reduced compared to unstratified PRS, but still greater than the GeRS for all outcomes except RheuArth.

### Estimating heritability explained by cis-heritable expression

AVENGEME estimated the SNP-based heritability of each phenotype based on PRS associations, with values ranging from 7.8% (95%CI=7.2%-8.4%) for CAD in UKB to 27.9% (95%CI=25.9%-30.0%) for Height in TEDS (Figure 7 and S13, Table S6). AVENGEME estimated the phenotypic variance explained by cis-heritable expression based on GeRS associations (GE-based heritability), returning estimates between 3.2% (95%CI=2.9%-3.5%) for Depression in UKB and 15.4% (95%CI=14.3%-16.6%) for IBD in UKB (Figure 7 and S13, Table S6). The proportion of SNP-based heritability explained by cis-heritable expression ranged from 26% for BMI in TEDS to 82% for RheuArth in UKB (Figure 7 and S13, Table S6). When restricting GeRS to colocalised features, the proportion of SNP-based heritability explained by cis-heritable expression ranged from 3% for ADHD in TEDS to 92% for RheuArth in UKB.

Estimates of the proportion of variants with no causal effect on the trait were broadly consistent when using PRS or GeRS, with PRS-based estimates ranging from 76.1% (95%CI=70.9%-80.6%) for GCSE in TEDS and 96.4% (95%CI=95.9%-96.9%) for IBD in UKB (Table S6).

## Discussion

This study has characterised the predictive utility of gene expression risk scores (GeRS), an approach that leverages gene expression summary statistics, GWAS summary statistics and target sample genotype data to infer genetic risk conferred via cis-regulated gene expression. We investigate factors affecting the predictive utility of GeRS, test whether GeRS can improve prediction over PRS alone, and estimate the proportion of SNP-based heritability that can be accounted for by cis-regulated expression. Our findings indicate GeRS represent a component of PRS, with GeRS explaining a substantial proportion of variance explained by PRS, suggesting GeRS may provide a useful approach for stratifying genetic risk into functional categories. Furthermore, this study finds GeRS generally provide small improvements in prediction over PRS alone, though GeRS can more substantial improvements in specific circumstances.

### Prediction using GeRS vs. PRS

GeRS typically explained less phenotypic variance than PRS derived using the same GWAS summary statistics. However, for several outcomes linear models combining GeRS and PRS did improve prediction over PRS alone. GeRS typically provided relative improvements of 1%-6% for the correlation between predicted and observed phenotype values, although for Rheumatoid Arthritis GeRS provided a 20.8% improvement when combined with pT+clump scores. All improvements in prediction provided by inclusion of GeRS were attenuated when using PRScs scores, which models LD as opposed to LD-based clumping, with GeRS only providing a 4% improvement for Rheumatoid Arthritis.

This pattern of results is likely due to the different method’s approaches and ability to jointly model variants in LD. The attenuated improvement for Rheumatoid Arthritis when using PRScs is particularly pronounced due to the methods ability to model effects within the MHC region as there are well-documented and strong HLA allele effects within the MHC region for Rheumatoid Arthritis (29). The PRScs method models all variation within the MHC region, where as pT+clump PRS considered only the strongest associated variant within the MHC region. In contrast, GeRS jointly models variants integrating their effect of gene expression, and then retains the single lead gene. This explanation is supported by our sensitivity analysis showing the gain in prediction for Rheumatoid Arthritis was also attenuated when compared to pT+clump PRS that were not clumped to a single variant within the MHC region. Nonetheless, the GeRS approach still provides some advantage over PRS in all cases, indicating that the functionally informed approach used by GeRS for jointly modelling variants better captures the risk in the MHC region than pT+clump or even PRScs can, possibly due to the documented eQTL effects in the locus altering expression of relevant HLA genes (30). Therefore, these results suggest GeRS can provide novel information over PRS alone, albeit in specific circumstances where multiple variants affecting gene expression are the causal risk factor. Given that the gene expression SNP-weights used in GeRS are derived using linear models, further improvement may be provided by using non-linear models that can capture haplotypes more effectively (31).

Inclusion of GeRS did not significantly improve prediction over PRS alone in the TEDS sample for any outcome. GeRS showed a similar correlation with Height and BMI as was found in the UK Biobank. These findings indicate the non-significant improvement in prediction when including GeRS is due to the smaller sample size of TEDS compared to UKB as this will reduce the power to detect small increases in prediction between models, and increase the likelihood of overfitting due to the large number of predictors in the model compared to the number of individuals in the sample. Approaches to efficiently integrate transcriptomic data without using many predictors would help alleviate this issue.

### Opportunities provided by GeRS

Although GeRS explain less variance than PRS, they may provide novel opportunities over PRS for several reasons. Firstly, GeRS are a gene-based genetic risk score, meaning the GeRS are well suited to stratification by biological pathways or other gene-based characteristics. Gene-based polygenic scores can also be created by restricting the variants considered to those proximal to genes (32). However, genetic variation proximal to a gene may have no effect on the gene’s expression or function. Secondly, GeRS are sensitive to the properties of the original gene expression dataset used to derive the expression SNP-weights and can capture gene expression within a range of contexts such as tissues and developmental stages. Therefore, GeRS could serve as a useful predictor for stratifying individuals based on the underlying aetiology of their disorder, addressing the possible that criticism of functionally agnostic PRS, that they are disconnected from aetiological considerations. Complex disorders are heterogenous at the phenotypic level and at the aetiological level. For example, it may be possible to stratify individuals based on the specific tissue underlying their condition.

### Factors affecting predictive utility of GeRS

Furthermore, models containing GeRS derived using multiple tissues improve prediction over the single best tissue, congruent with a previous study (12).

We found the relative predictive utility of GeRS derived using different SNP-weight sets was strongly correlated with the number of genes captured by the SNP-weight set. This is likely due to a multitude of factors including the sample size and quality of the original gene expression dataset. Both of these factors will increase the number of genes captured by the SNP-weight set by detecting more genes with significantly heritable cis-regulated expression, and increase the variance in gene expression the SNP-weights explain out-of-sample. It is likely that the relevance of the tissue to the outcome is also an important factor influencing the predictive utility of an outcome, however the sample size and number of features have a larger effect on the predictive utility of GeRS due to the moderately correlated cis-regulated expression across tissues enabling tissues irrelevant to the outcome to act as a proxy for gene expression within relevant but unavailable tissues.

### Quantifying heritability accounted for by cis-regulated expression

GeRS capture only a small amount of novel phenotypic variance compared to PRS, indicating that GeRS largely capture a component of risk captured by PRS. These findings are congruent with a previous study modelling the genome and genetically regulated transcriptome using CORE GREML (14). We estimate the proportion of phenotypic variance that can be explained by *cis*-regulated gene expression and compare the results to SNP-based heritability estimates using PRS results. Across the phenotypes we estimate *cis*-regulated gene expression explains 26%-82% of SNP-based heritability. However, due to linkage disequilibrium GeRS are likely to capture effects mediated through other mechanisms. To more accurately estimate the proportion of SNP-based heritability accounted for by *cis*-regulated expression, we restricted the analysis to genes that colocalise and their association is therefore unlikely to be driven by linkage. When restricting the analysis to colocalised genes, we estimated 3% to 92% of SNP-based heritability was accounted for by *cis*-regulated expression. For most outcomes this supports previous research showing strong enrichment of eQTLs in GWAS summary statistics (5, 6). Our findings suggest that restricting GeRS to colocalised genes will reduce their predictive utility but may provide a more accurate estimate of an individual’s risk mediated via *cis*-regulated expression. This raises a further issue for gene-based polygenic scores, as they are more liable to capturing linkage effects and there is no option to restrict analyses to colocalised genes. Even when restricting our analysis to colocalised genes, our estimates of phenotypic variance attributable to *cis*-regulated expression may still be upwardly biased due to GeRS capturing effects driven by horizontal pleiotropy as opposed to vertical pleiotropy (mediation). An example of horizontal pleiotropy would be where a disease-associated variant is an eQTL for a gene, but the variant confers risk for the disease via another mechanistic route, such as trans eQTL effects. A recently developed method called MESC can be used to identify the variance explained by vertical pleiotropy (mediation) alone (33). Indeed, the results reported by MESC are lower than the estimates based on GeRS in this study.

### Opportunities for GeRS based on observed expression

Although the colocalization and tissue specificity of genes did not improve prediction of GeRS when based on *predicted* expression in the target sample, restricting genes by these criteria is likely to improve the predictive utility of GeRS derived using *observed* gene expression in the target sample. This is supported by a previous study which found GeRS derived using GWAS summary statistics and eQTL data, and *observed* gene expression data, could substantially improve prediction over PRS but only when using eQTL data from the relevant tissue and restricting the risk scores to colocalised genes (23). Tissue specificity and colocalization is more important when integrating with observed gene expression as the GeRS must capture genuine differences in expression associated with the outcome. Future research exploring the predictive utility of GeRS derived using TWAS results and observed expression is warranted.

In summary, this study has demonstrated that GeRS explain a substantial proportion of variance for a range of outcomes, with multiple tissue GeRS explaining more variance than the single best tissue. Furthermore, we demonstrate that GeRS can improve prediction of outcomes over PRS alone in specific circumstances, where multiple eQTL effects must be considered to fully capture the genetic risk conferred by a locus. However, the results largely indicate that GeRS capture a component of risk captured by functionally agnostic PRS, and estimates of variance explained by cis-regulated expression is 26%-82% of total SNP-based heritability, although these estimates likely captures risk not only mediated via cis-regulated expression due to horizontal pleiotropy and linkage. In conclusion, GeRS may serve as a useful research tool by providing a novel opportunity to stratify genetic risk by expression within specific tissues, developmental stage, and other gene-based characteristics.

## URLS

- LDSC HapMap 3 SNP-list: https://data.broadinstitute.org/alkesgroup/LDSCORE/w_hm3.snplist.bz2
- LDSC Munge Sumstats: https://github.com/bulik/ldsc/blob/master/munge_sumstats.py
- Impute.me: https://impute.me/
- GenoPred website: https://opain.github.io/GenoPred

## Acknowledgements

We thank Paul O’Reilly, Sam Choi and Alexander Gusev for useful discussion.

This paper represents independent research funded by the UK Medical Research Council (MR/N015746/1 and MR/S0151132), and the National Institute for Health Research (NIHR) Biomedical Research Centre at South London and Maudsley NHS Foundation Trust and King’s College London. AF is funded by the National Institute of Health grant R01AG054628. The authors acknowledge use of the research computing facility at King’s College London, Rosalind (https://rosalind.kcl.ac.uk), which is delivered in partnership with the NIHR Maudsley BRC, and part-funded by capital equipment grants from the Maudsley Charity (award 980) and Guy’s & St. Thomas’ Charity (TR130505). The views expressed are those of the authors and not necessarily those of the NHS, the NIHR or the Department of Health and Social Care. We thank the research participants and employees of 23andMe for making the work regarding Depression possible.

UKB: This research was conducted under UK Biobank application 18177.

TEDS: We gratefully acknowledge the ongoing contribution of the participants in TEDS and their families. TEDS is supported by UK Medical Research Council Program Grant MR/M021475/1 (and previously Grant G0901245) (to Robert Plomin (R.P).), with additional support from National Institutes of Health Grant AG046938. The research leading to these results has also received funding from the European Research Council under the European Union’s Seventh Framework Programme (FP7/2007-2013) ERC Grant Agreement 295366. R.P. is supported by Medical Research Council Research Professor Award G19/2. KR is supported by a Sir Henry Wellcome Postdoctoral Fellowship.

