## Supplementary Information for "Imputed Gene Expression Risk Scores: A Functionally Informed Component of Polygenic Risk"

### UKB Outcome definitions

*Depression.* UKB participants were coded as depression cases if they met the Composite International Diagnostic Interview Short Form criteria for lifetime depression which was assessed in the online Mental Health Questionnaire (MHQ) using scoring protocols proposed by Davis et al (1). Depression cases were screened for indications of schizophrenia or bipolar disorder according to the MHQ. Controls excluded if they show any psychiatric indications according to the MHQ or depression indications according to: ICD-10 diagnoses; endorsement of self-reported depression; endorsement of current antidepressant usage; single or current depression according to the criteria adopted by Smith, et al (2). Further details of the exclusion criteria have been previously described (3).

*T2D.* Cases were identified based on a combination of hospital episode statistics, using both ICD-9 and ICD-10, the national death register, and self-reported questionnaire data. In order to classify as a case for type 2 diabetes, self-reported type 2 or generic diabetes status was established in the nurse interview and the touchscreen questionnaire. However, participants were only classified as cases when they reported in the questionnaire that they had not been treated with insulin in the first year after diagnosis and had been diagnosed after the age of 35 years. Type 2 diabetes controls did not fulfil these criteria and did not have any other types of diabetes. Further details of the T2D definition have been previously published (4).

*Coronary artery disease (CAD).* Participants who were registered in the hospital in-patient data or the death register to have had ischemic heart diseases, or participants who had coronary revascularization operations were classified as coronary artery disease cases in this study. If participants self-reported those conditions in the nurse interview or the touchscreen questionnaire, they were also considered to have coronary artery disease. Coronary artery disease controls did not fulfil those criteria. Further details of the CAD definition have been previously published (4).

*Autoimmune diseases (IBD, RheuArth):* UKB participants were coded as autoimmune cases if at least two of the following measures were observed: ICD-10 diagnoses from Hospital Episode Statistics; endorsement of self-reported autoimmune diseases; endorsement of prescription medication for the corresponding autoimmune diseases. More than one hospital admission for the respective autoimmune conditions was also sufficient. Controls were excluded if any of the following were observed: Pernicious Anemia, Autoimmune Thyroid Disease, Type 1 diabetes, Multiple Sclerosis, Myasthenia Gravis, Coeliac, Inflammatory Bowel Disease, Hidradenitis Suppurativa, Pemphigoid/Pemphigus, Psoriasis, Ankylosing Spondylitis, Polymyalgia Rheumatica/Giant Cell Arteritis, Psoriatic Arthritis, Rheumatoid Arthritis, Sjögren Syndrome, Systemic Lupus Erythematosus.

*Intelligence* was defined using the Fluid intelligence score variable. Fluid intelligence was assessed using the 13 item UKB Touch-screen Fluid intelligence test (5). The test measures the capacity to solve problems that require logic and reasoning ability, independent of acquired knowledge. The fluid intelligence variable was derived by UKB as an unweighted sum of the number of correct answers, assigning a score of 0 to unanswered questions.

*Height* was defined using the Standing height variable (Field ID: f.50.0.0).

*BMI* was defined using the Body mass index variable (Field ID: f.21001.0.0).

### TEDS outcome definitions

*Height*: Self-reported height was assessed at age 21.

*BMI*: BMI was calculated as self-reported weight in kilograms divided by height in meters squared (kg/m^2^) at age 21.

*Educational Achievement*: Results for standardized tests taken at the end of compulsory education in the United Kingdom (General Certificate of Secondary Education; GCSE) were obtained for twins at mean age 16.3 years (SD = 0.29) via self-report or parent-report. Grades were coded from 4 (G; the minimum pass grade) to 11 (A*; the highest possible grade), with the U fail grade coded as missing. A composite score was calculated as the arithmetic mean of the compulsory core subjects—Maths, English, and Science. Further information on this definition of Educational Attainment has been previously published (6).

*Attention-Deficit Hyperactivity Disorder (ADHD) Symptoms*: At age 11.5 (SD = 0.69) and 16.3 (SD = 0.69), parents reported on twins’ ADHD symptoms via the Strength and Difficulties Questionnaire (7) hyperactivity subscale (three-point Likert scale) and the Conners’ rating scales (CPRS-R; four-point Likert scale) (8) on hyperactivity and inattention. A composite score was created as the arithmetic mean of the sex and age z-standardized scales. Where ratings were available at one assessment only, this value was used to maximize sample size.

### LDSC munge_sumstats.py settings used

- Removes SNPs not in HapMap3 list (w_hm3.snplist)
- Removes SNPs with missing values.
- Removes SNPs with INFO <= 0.6.
- Removes SNPs with MAF <= 0.01.
- Removes SNPs with out-of-bounds p-values.
- Removes variants that were not SNPs or were strand-ambiguous.
- Removes SNPs with duplicated RS numbers.
- Removes SNPs with N < (90th percentile N) / 2.
- Removes SNPs whose alleles did not match HapMap3 list (w_hm3.snplist)

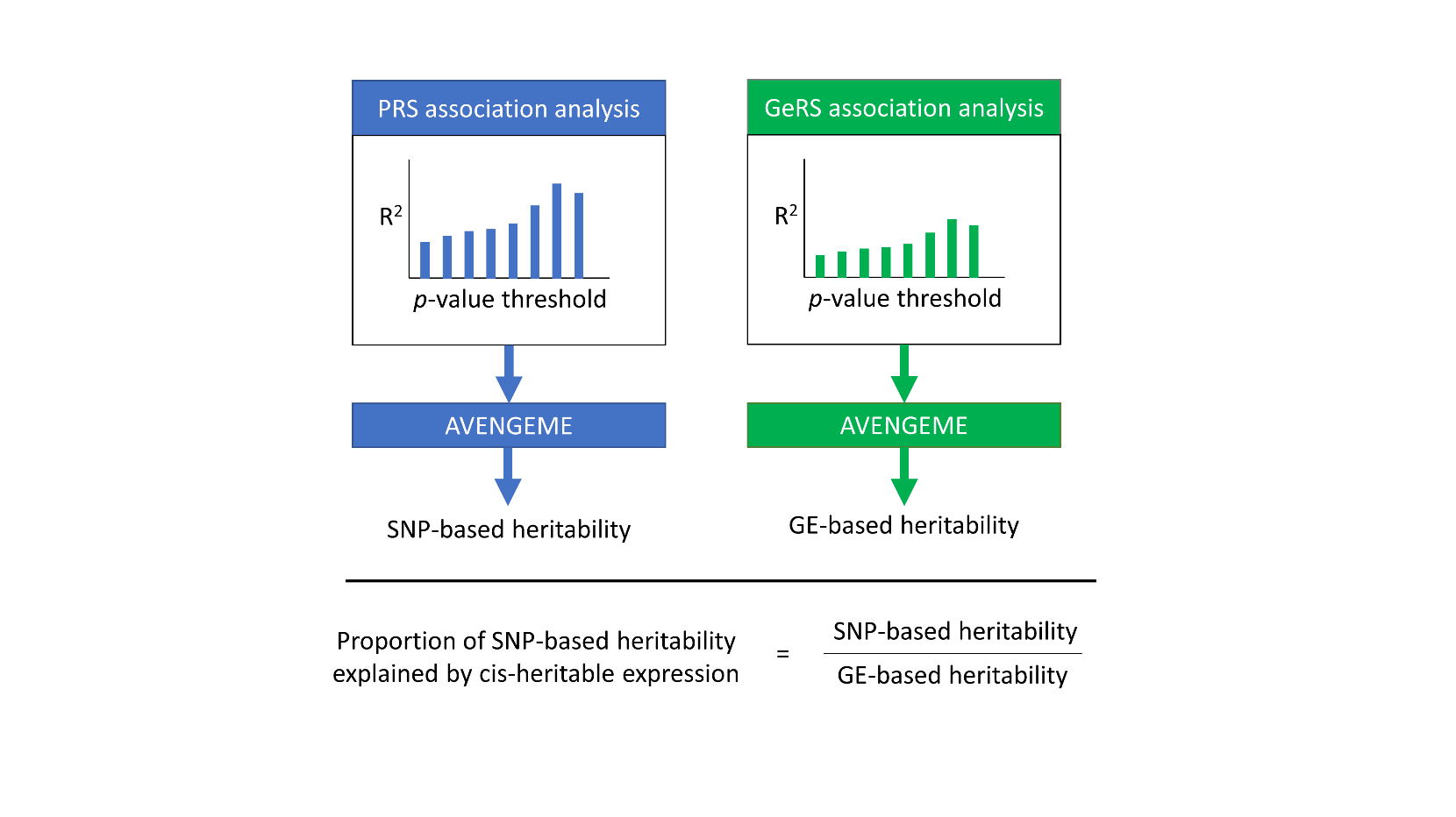

Supplementary Figure 1. Schematic representation of analysis estimating the proportion of SNP-based heritability explained by cis-heritable expression.

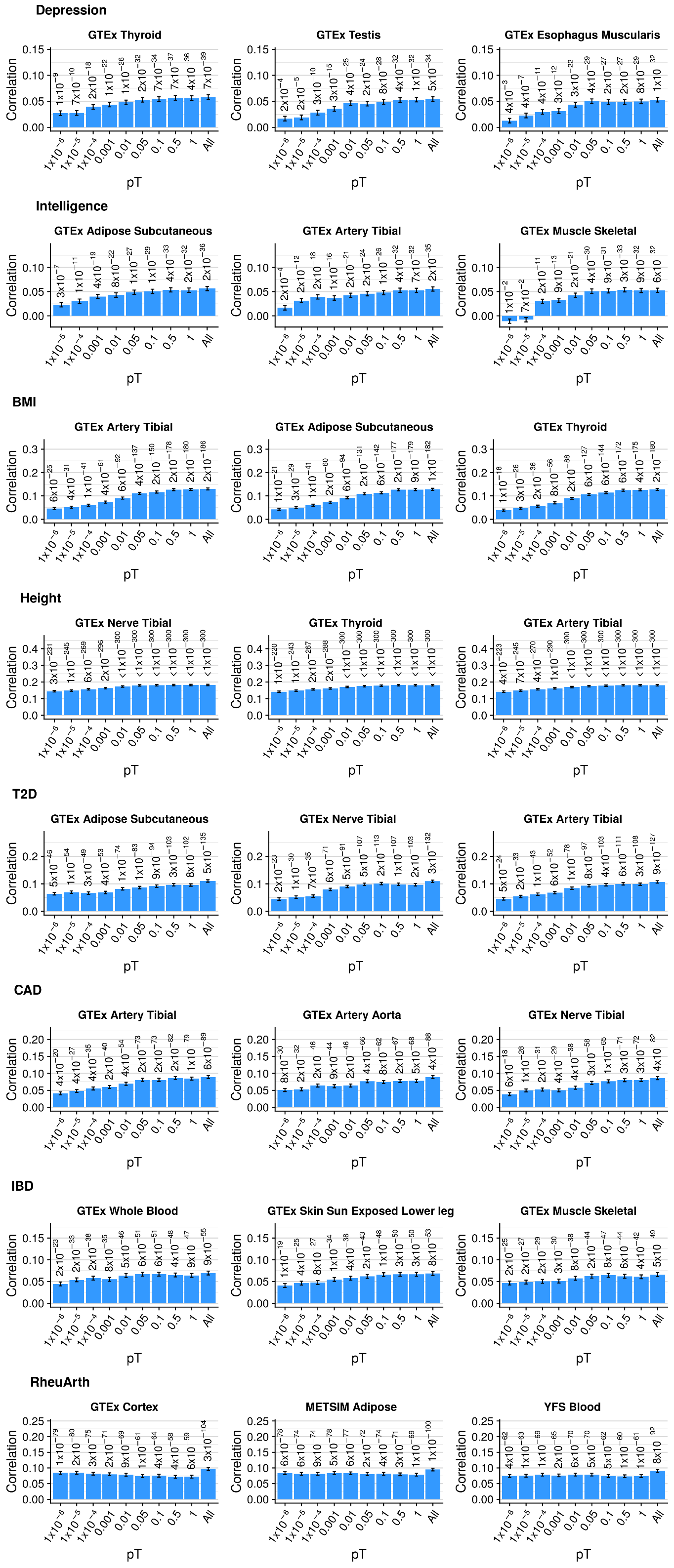

Supplementary Figure 2 part 1. Correlation between GeRS for three outcomes in UKB across p-value thresholds. The all bar indicates the correlation between observed and predicted values from an elastic net model including all p-value thresholds. Error bars represent the standard error of the correlation. GeRS are based on a single SNP-weight sets. Figure only shows results for the three SNP-weights with the strongest correlation between predicted and observed values. Values above bars are p-values indicating whether the correlation is significantly different from zero.

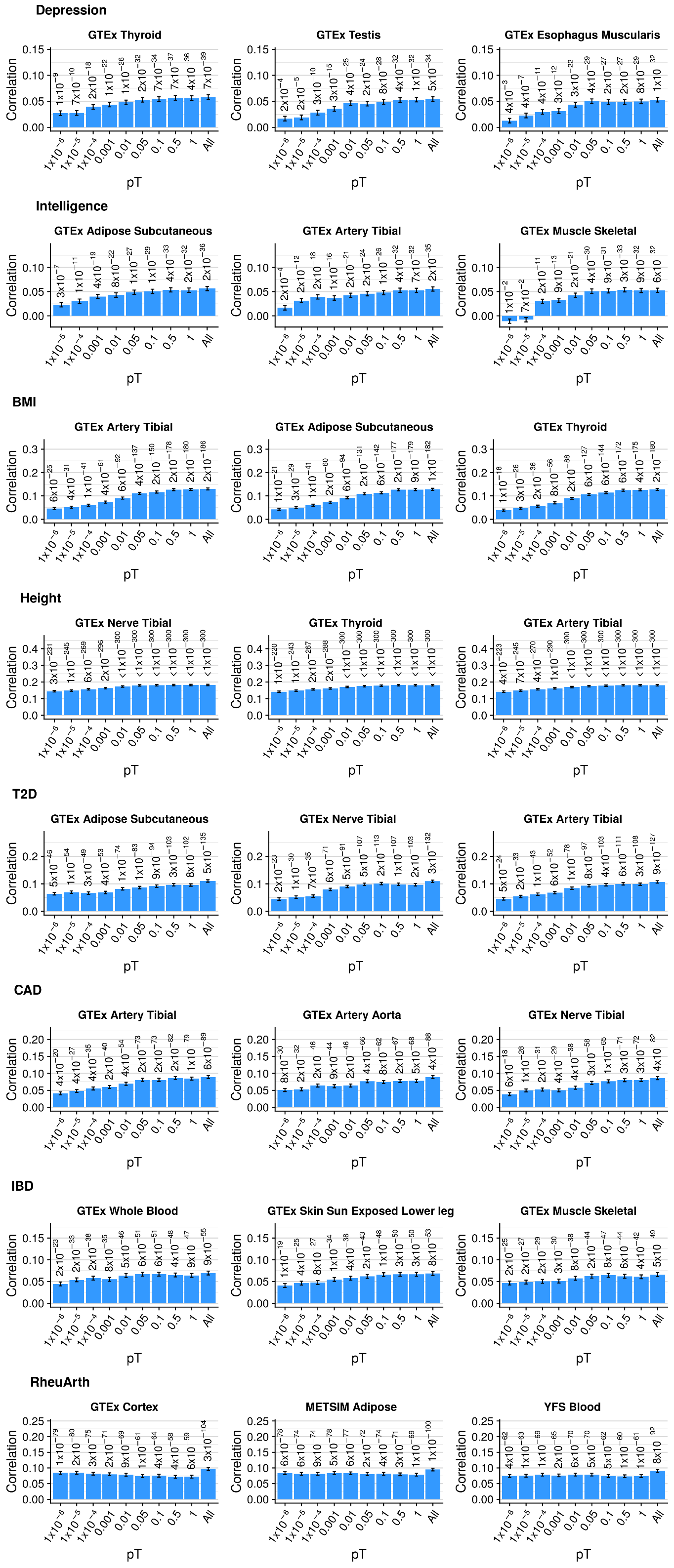

Supplementary Figure 2 part 2. Correlation between GeRS for three outcomes in UKB across p-value thresholds. The all bar indicates the correlation between observed and predicted values from an elastic net model including all p-value thresholds. Error bars represent the standard error of the correlation. GeRS are based on a single SNP-weight sets. Figure only shows results for the three SNP-weights with the strongest correlation between predicted and observed values. Values above bars are p-values indicating whether the correlation is significantly different from zero.

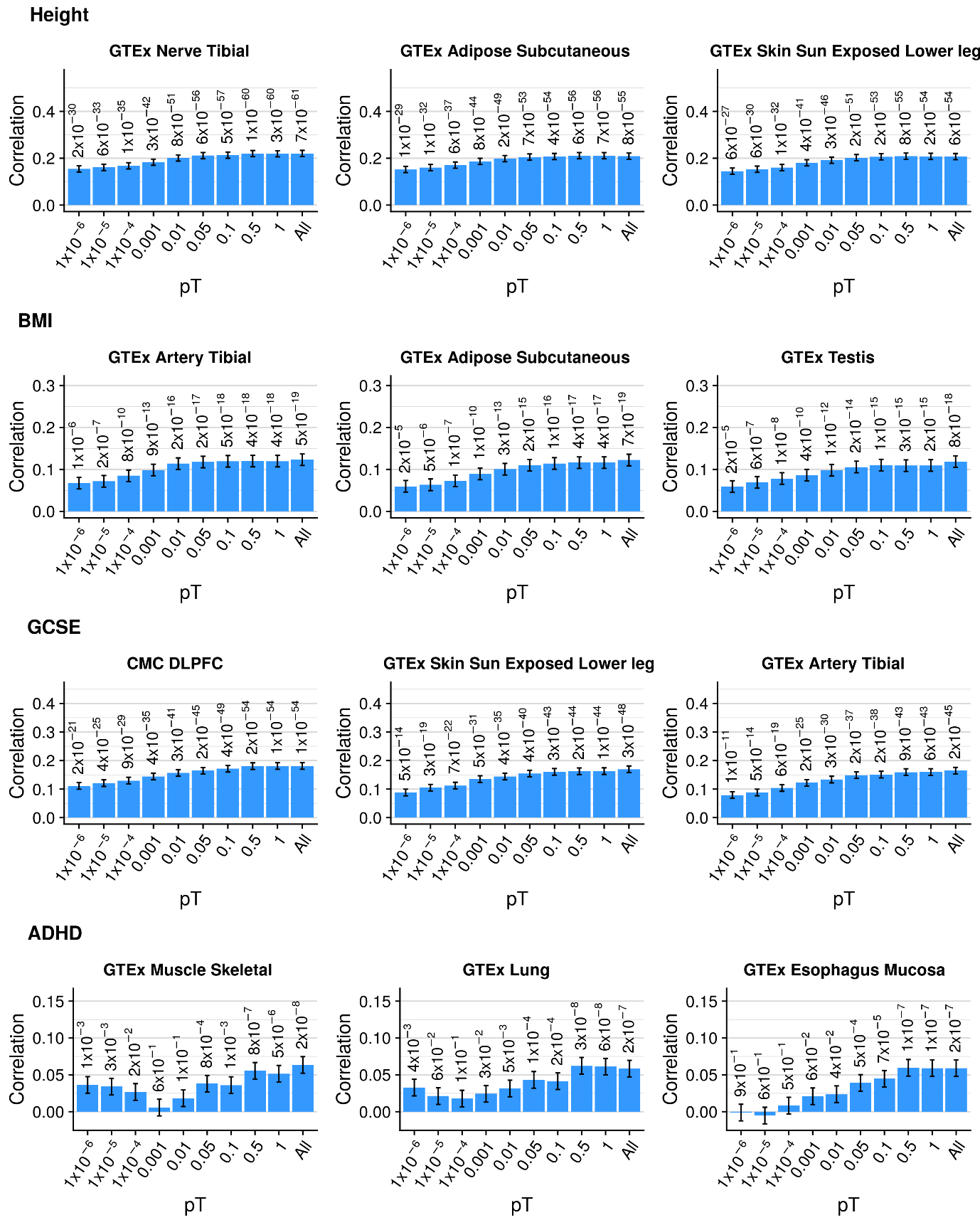

Supplementary Figure 3. Correlation between GeRS for three outcomes in TEDS across p-value thresholds. The all bar indicates the correlation between observed and predicted values from an elastic net model including all p-value thresholds. Error bars represent the standard error of the correlation. GeRS are based on a single SNP-weight sets. Figure only shows results for the three SNP-weights with the strongest correlation between predicted and observed values. Values above bars are p-values indicating whether the correlation is significantly different from zero.

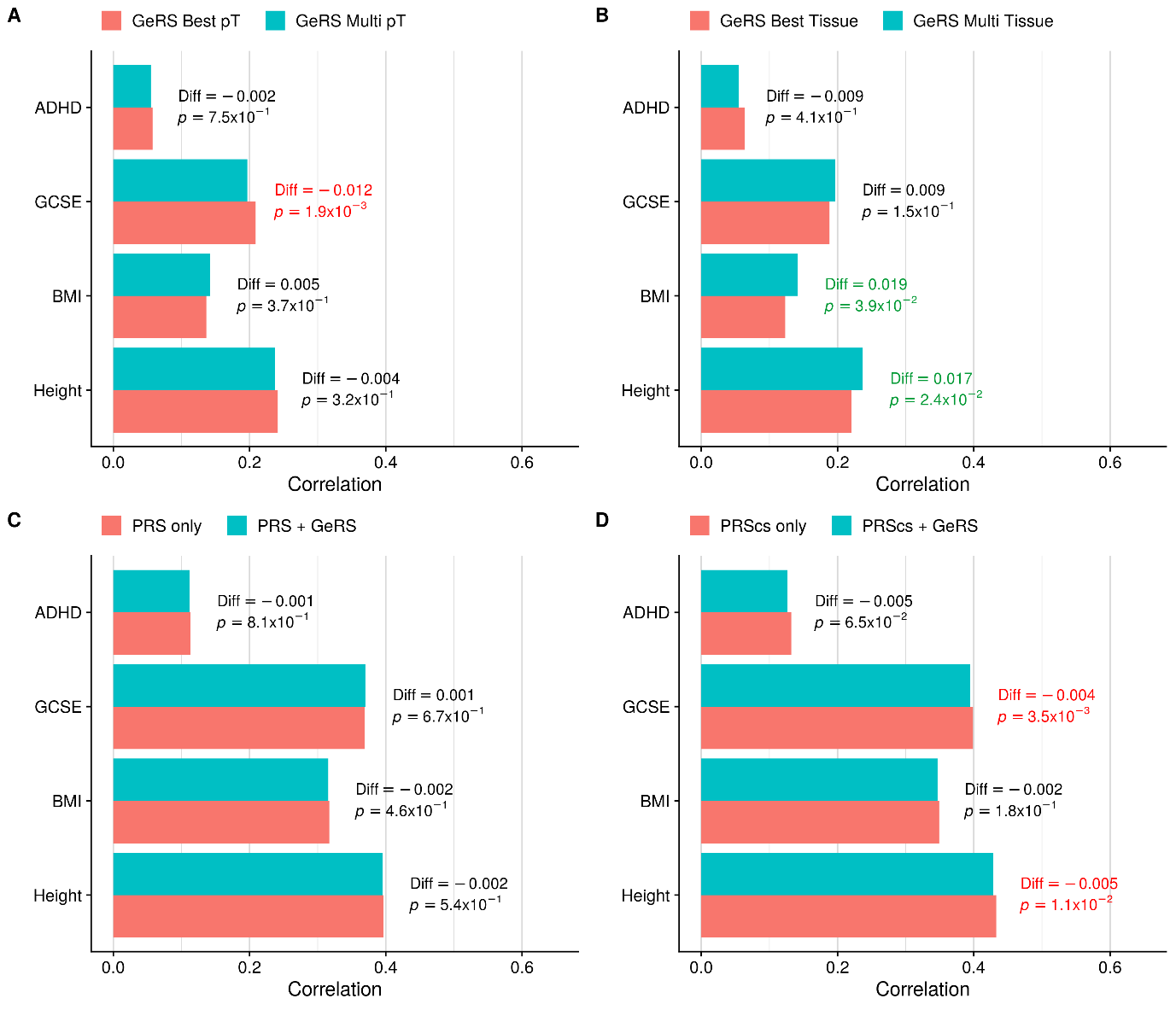

Supplementary Figure 4. Comparing the predictive utility of GeRS and PRS in TEDS. A) Compares the predictive utility of models containing GeRS across SNP-weight sets based on the single best p-value threshold and models containing GeRS across all p-value thresholds. B) Compares the predictive utility of models containing GeRS across p-value thresholds based on the single best SNP-weight set and models containing GeRS based on all SNP-weight sets. C) Compares the predictive utility of models containing PRS and models containing GeRS and PRS. D) Compares the predictive utility of models containing models also containing PRS derived using PRScs, and models also containing GeRS. Values on the right of each bar indicate the absolute difference in predicted-observed correlation between the full and nested model. Values on the right are coloured in green to indicate a significant increase in prediction, and red to indicate a significant decrease in prediction.

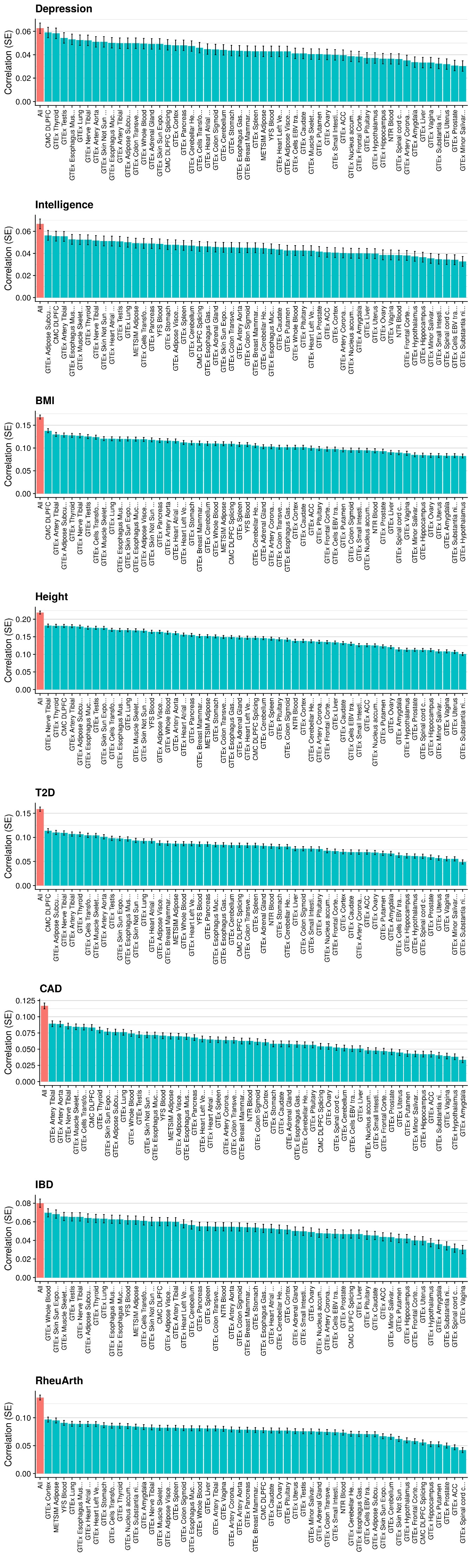

Supplementary Figure 5 part 1. Predictive utility of GeRS in UKB derived from each SNP-weight set, and a model combining all SNP-weight sets. The red bar indicates the model containing all SNP-weight sets.

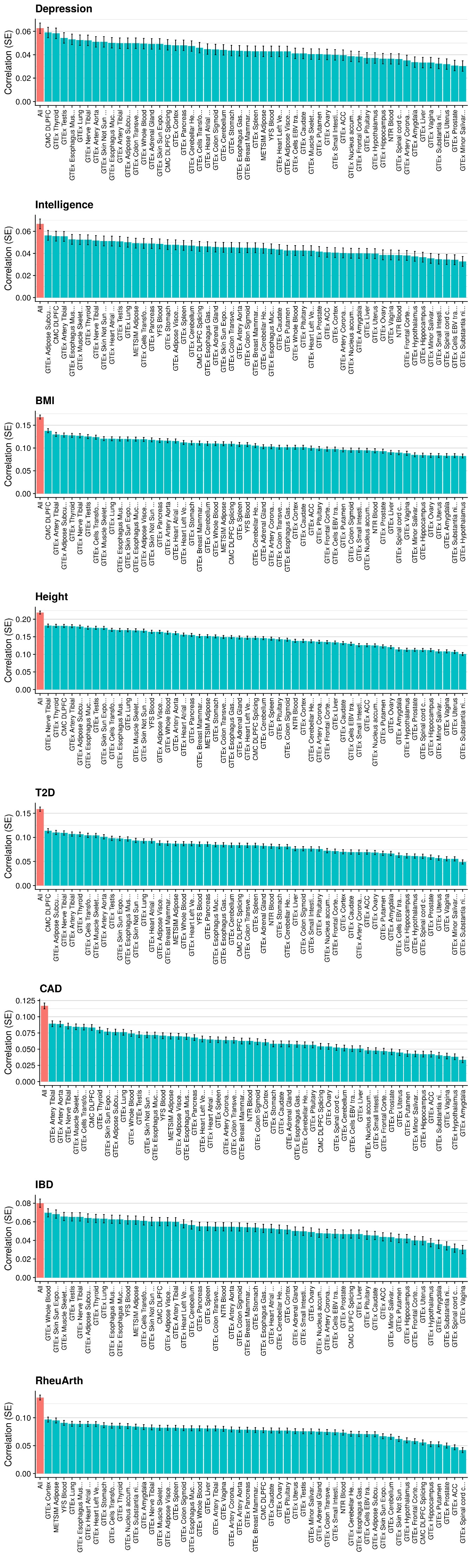

Supplementary Figure 4 part 2. Predictive utility of GeRS in UKB derived from each SNP-weight set, and a model combining all SNP-weight sets. The red bar indicates the model containing all SNP-weight sets.

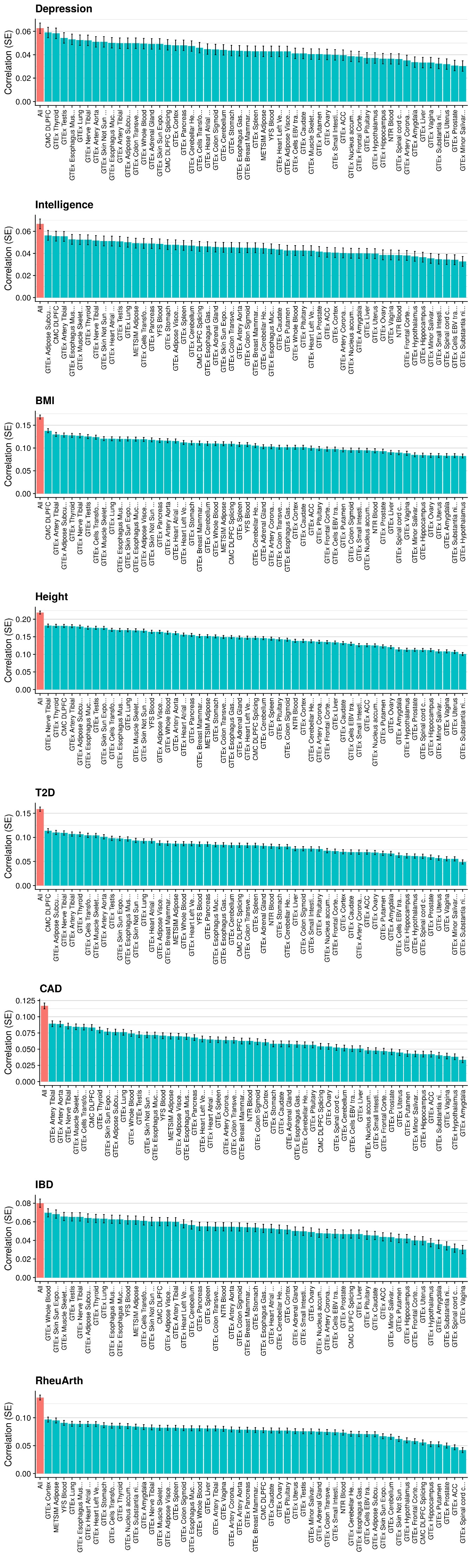

Supplementary Figure 4 part 3. Predictive utility of GeRS in UKB derived from each SNP-weight set, and a model combining all SNP-weight sets. The red bar indicates the model containing all SNP-weight sets.

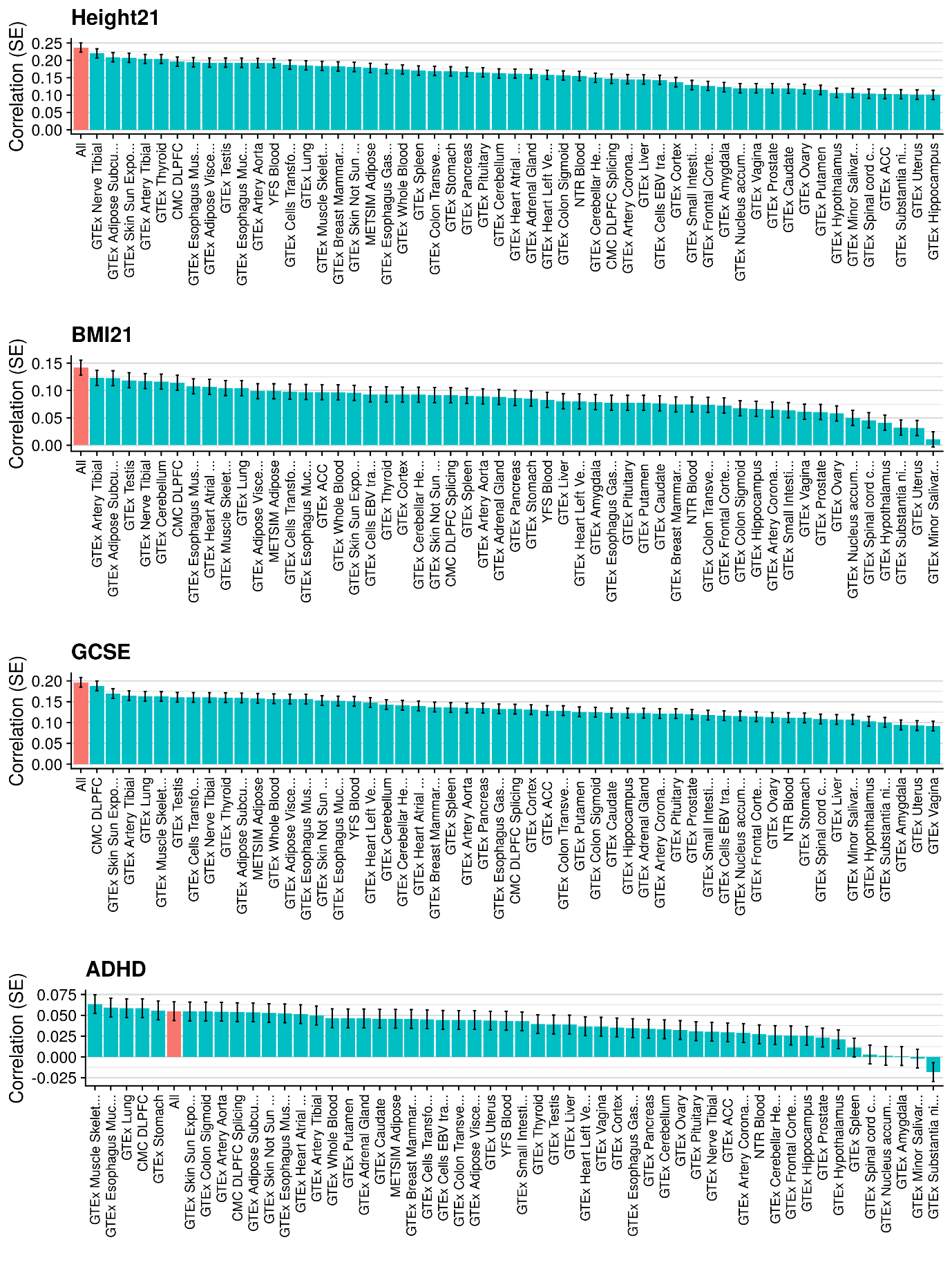

Supplementary Figure 6. Predictive utility of GeRS in TEDS derived from each SNP-weight set, and a model combining all SNP-weight sets. The red bar indicates the model containing all SNP-weight sets.

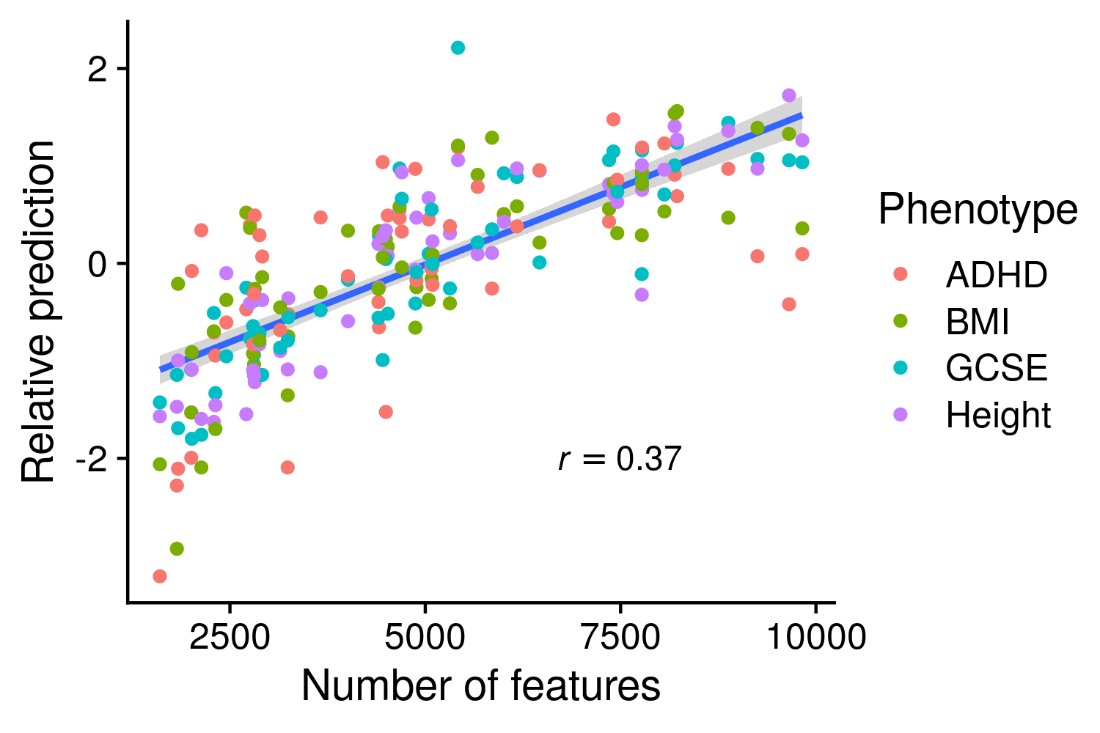

Supplementary Figure 7. Relationship between the predictive utility of GeRS and the number of features within each SNP-weight set in TEDS. The y-axis shows the correlation between observed and predicted values, standardised within each outcome.

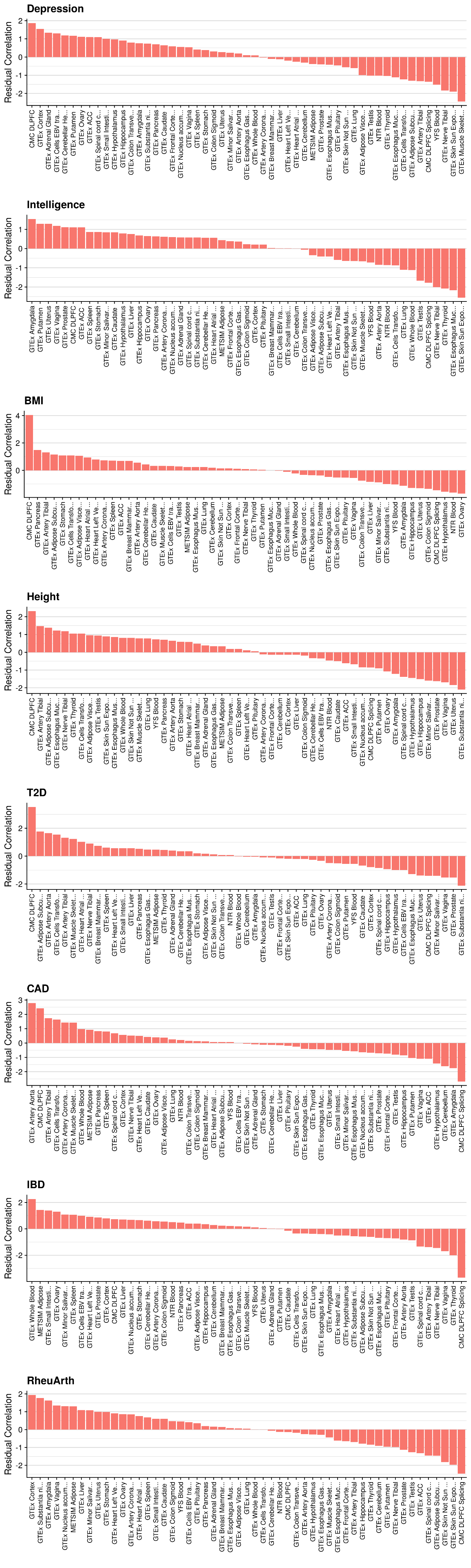

Supplementary Figure 8 part 1. Relative predictive utility of GeRS for each SNP-weight set in UKB after adjusting for the number of features within each SNP-weight set.

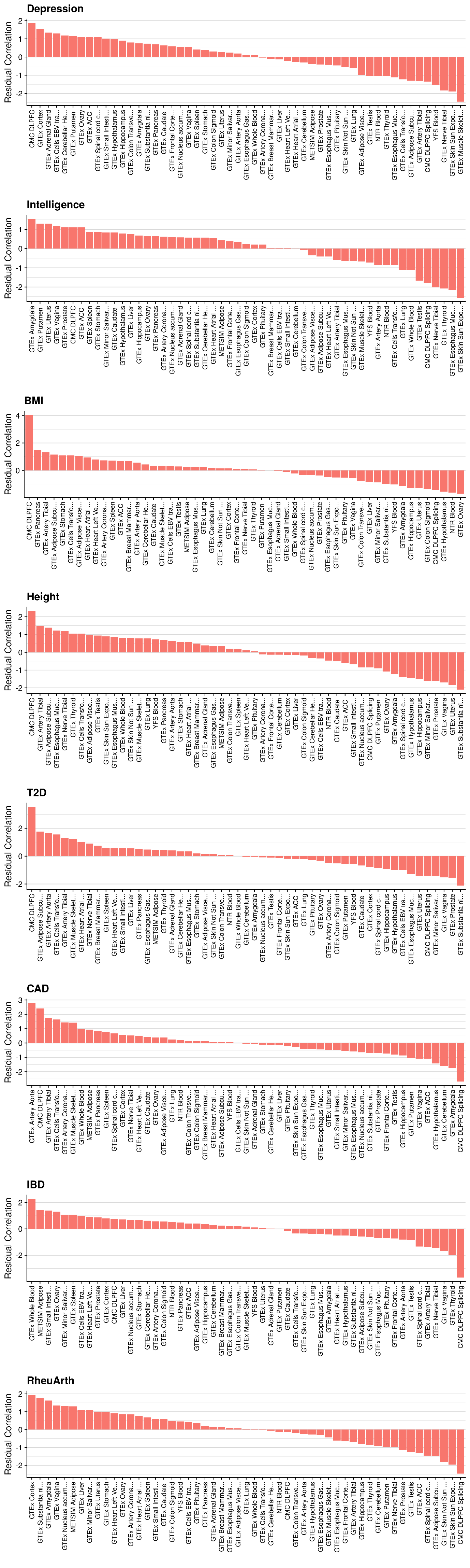

Supplementary Figure 8 part 2. Relative predictive utility of GeRS for each SNP-weight set in UKB after adjusting for the number of features within each SNP-weight set.

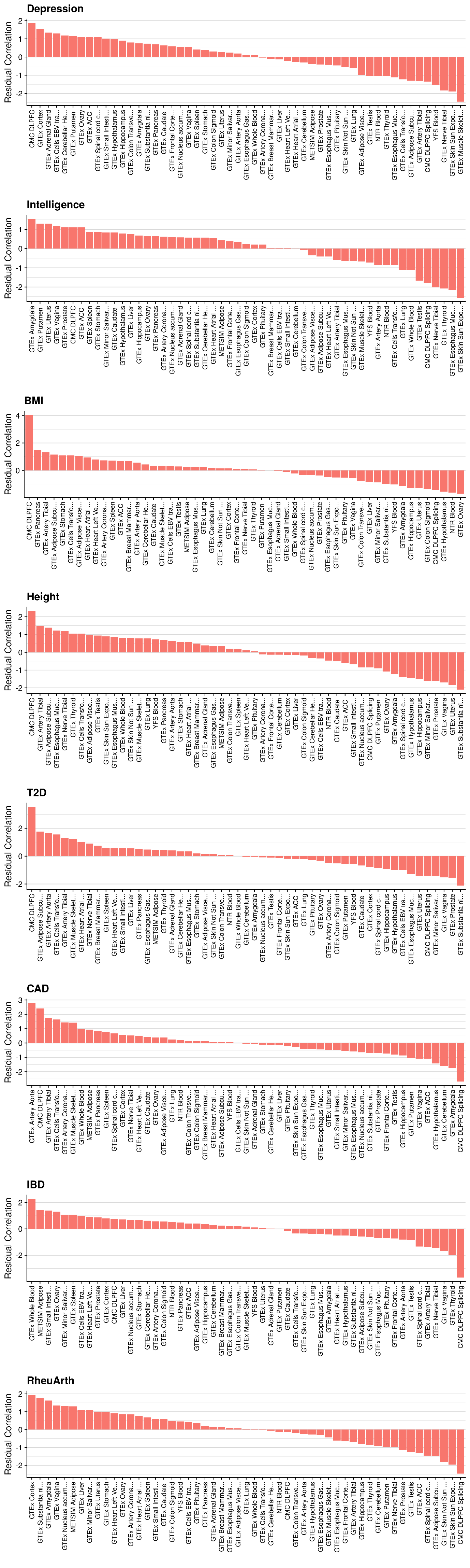

Supplementary Figure 8 part 3. Relative predictive utility of GeRS for each SNP-weight set in UKB after adjusting for the number of features within each SNP-weight set.

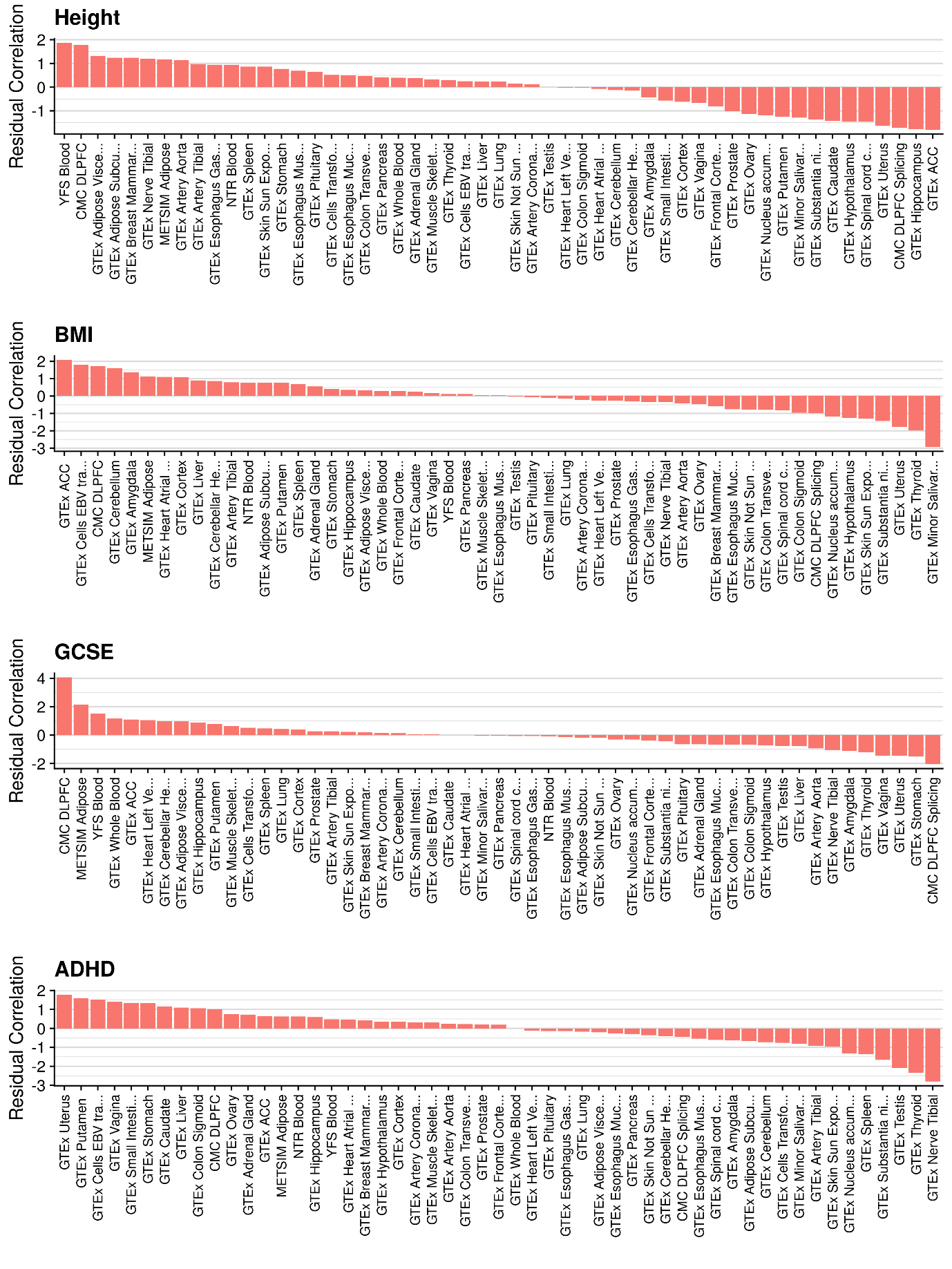

Supplementary Figure 9. Relative predictive utility of GeRS for each SNP-weight set in TEDS after adjusting for the number of features within each SNP-weight set.

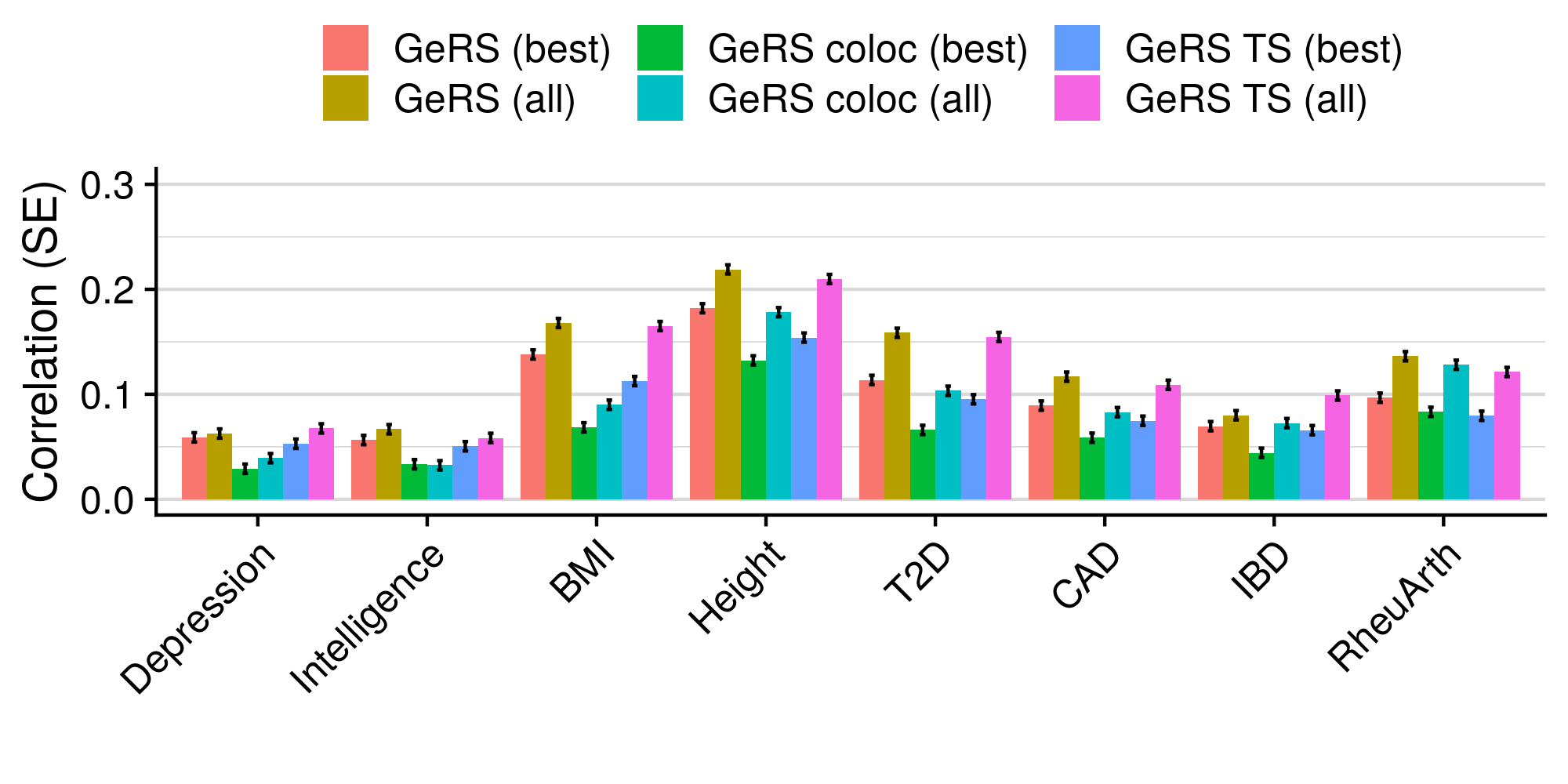

Supplementary Figure 10. Predictive utility of GeRS in UKB, using all features (GeRS), using colocalized features (PP4 > 0.8) (GeRS coloc), and using tissue-specific features (GeRS TS). The predictive utility for the single best SNP-weight set (best) and all SNP-weight sets combined (all) are shown.

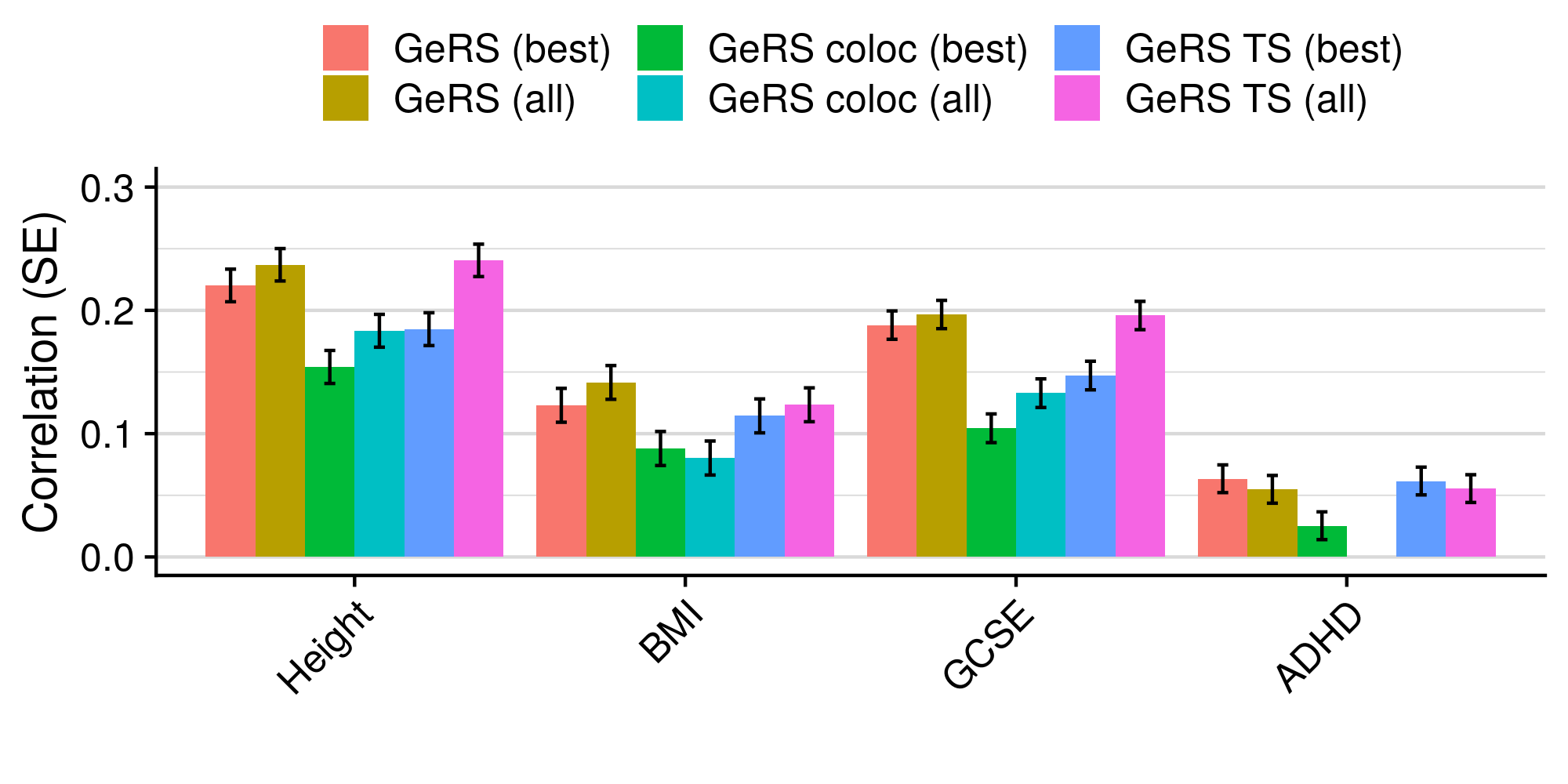

Supplementary Figure 11. Predictive utility of GeRS in TEDS, using all features (GeRS), using colocalized features (PP4 > 0.8) (GeRS coloc), and using tissue-specific features (GeRS TS). The predictive utility for the single best SNP-weight set (best) and all SNP-weight sets combined (all) are shown.

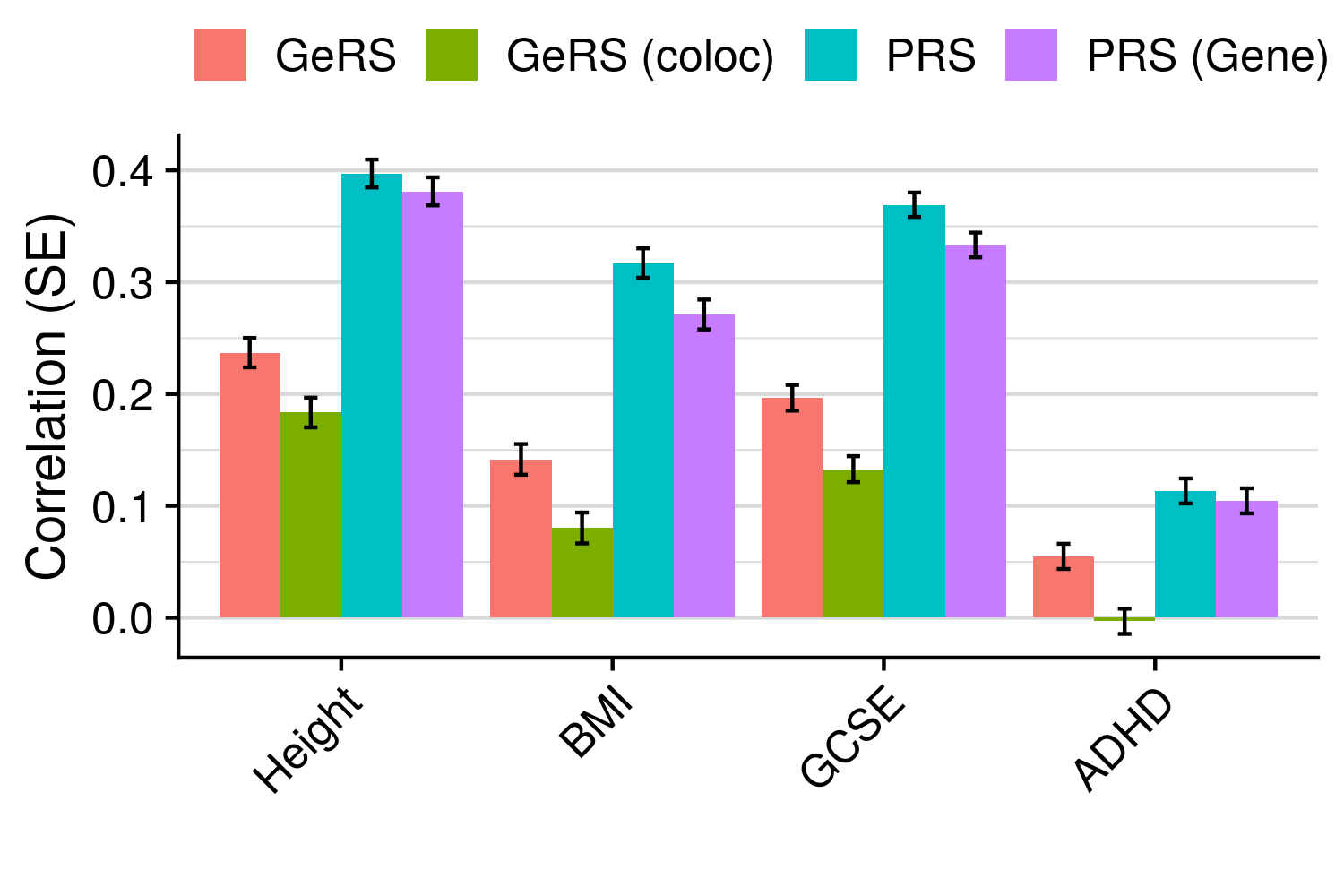

Supplementary Figure 12. Shows the correlation between predicted and observed values in TEDS for models. GeRS = All SNP-weight set GeRS; GeRS (coloc) = All SNP-weight set GeRS restricted to genes with a colocalization PP4 > 0.8; PRS = Genome-wide PRS; PRS (Gene) = PRS restricted to gene regions considered by GeRS.

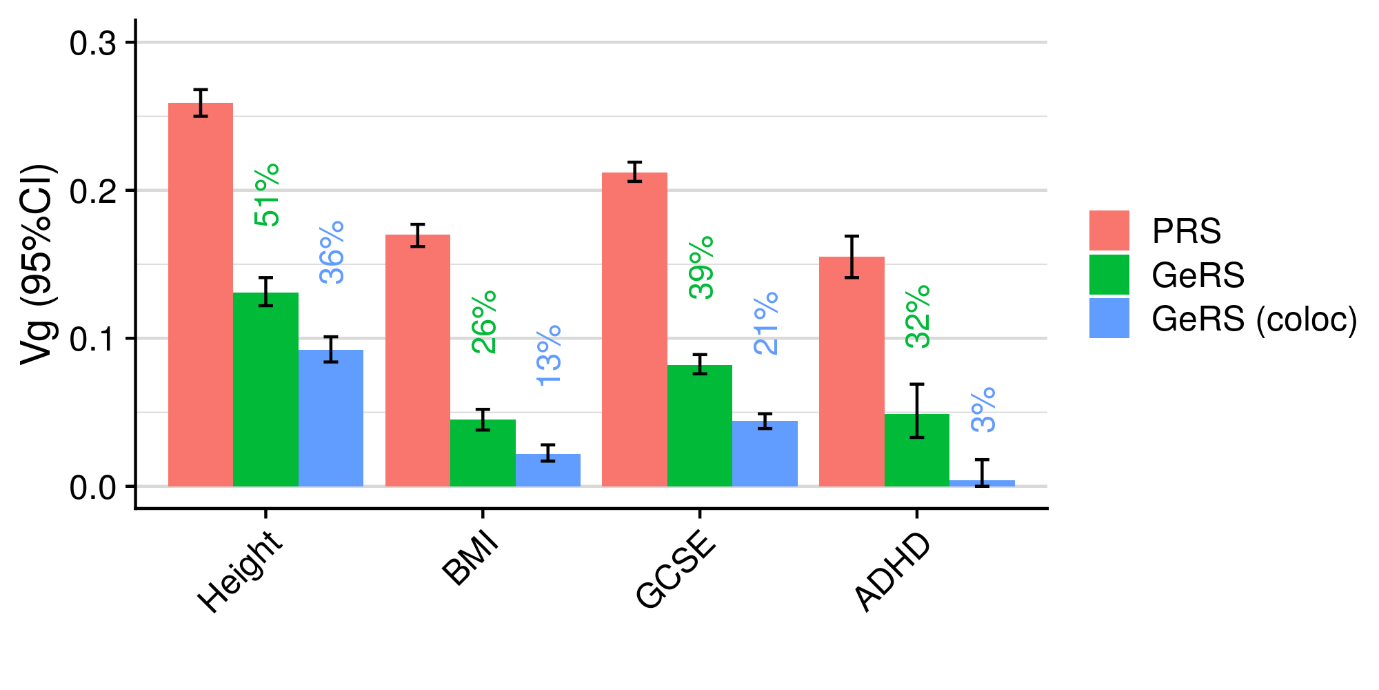

Supplementary Figure 13. Estimates of SNP-based heritability and GE-based heritability for outcomes in TEDS. PRS indicates the SNP-heritability as estimated using PRS association results in AVENGEME. GeRS indicates the GE-based heritability as estimated using GeRS association results in AVENGEME. The value above each bar indicates the proportion of SNP-based heritability accounted for by cis-regulated expression (Green=GeRS/PRS, Blue=GeRS (coloc)/PRS).
